# Cooperative engagement and subsequent selective displacement of SR proteins define the pre-mRNA 3D structural scaffold for early spliceosome assembly

**DOI:** 10.1101/2021.12.01.470860

**Authors:** Kaushik Saha, Gourisankar Ghosh

**Affiliations:** Department of Chemistry and Biochemistry, University of California San Diego, 9500 Gilman Drive, La Jolla, CA 92093-0375

## Abstract

We recently reported that serine-arginine-rich (SR) protein-mediated pre-mRNA structural remodeling generates a pre-mRNA 3D structural scaffold that is stably recognized by the early spliceosomal components. However, the intermediate steps between the free pre-mRNA and the assembled early spliceosome are not yet characterized. By probing the early spliceosomal complexes *in vitro* and RNA-protein interactions *in vivo*, we show that the SR proteins bind the pre-mRNAs cooperatively generating a substrate that recruits U1 snRNP and U2AF65 in a splice signal-independent manner. Excess U1 snRNP selectively displaces some of the SR protein molecules from the pre-mRNA generating the substrate for splice signal-specific, sequential recognition by U1 snRNP, U2AF65, and U2AF35. Our work thus identifies a novel function of U1 snRNP in mammalian splicing substrate definition, explains the need for excess U1 snRNP compared to other U snRNPs *in vivo*, demonstrates how excess SR proteins could inhibit splicing, and provides a conceptual basis to examine if this mechanism of splicing substrate definition is employed by other splicing regulatory proteins.

## INTRODUCTION

The early spliceosome defines the mammalian splicing substrate by recognizing four major splice signals, namely 5′ and 3′ splice sites (SS) located at the 5′- and 3′-ends of the intron as well as branch-point sites (BS) and polypyrimidine tracts (PPT) near the 3′SS. These four splice signals are recognized by U1 small nuclear ribonucleoprotein (U1 snRNP), U2 auxiliary factor 35 (U2AF35), splicing factor 1 (SF1), and U2 auxiliary factor 65 (U2AF65), respectively (1). U2AF65 is considered to be the primary factor that defines the 3′SS (2, 3). However, these recognition events are not governed solely by the splice signal sequences but by a combination of features embedded within the pre-mRNA strand, collectively known as the ‘splicing code’ (4). The features of a pre-mRNA splicing code may include the sequence of the splice signals, the pre-mRNA secondary structure (5, 6), the splicing regulatory elements (SREs) (7), and the pre-mRNA 3D structural scaffold (8). The four aspects of the splicing code could also mutually regulate each other: secondary structure could control the exposure, occlusion (6), and proximity (9) of SREs and splice signals, pre-mRNA 3D structural scaffold could regulate the functionality of SREs, RNA binding proteins (RBPs) recruited by the SREs could modulate the pre-mRNA scaffold, and/or the secondary structure, and mutation in splice signal sequences could disrupt the pre-mRNA 3D scaffold (8).

Serine-arginine-rich (SR) proteins are key RBPs in the regulation of early spliceosome assembly, and deregulation of this process causes widespread splicing changes leading to a wide range of diseases (10–18). Accurate correlation of binding of RBPs to the cognate SREs within the pre-mRNA with the splicing outcome is critical for elucidating the mammalian ‘splicing code’ (19). Currently, several mechanistic aspects of the functions of SR proteins remain unclear. Firstly, it is recently reported that the 3D scaffold of a pre-mRNA can be recognized by the early spliceosomal components and structural remodeling of the scaffold by SR proteins enhances the stability of the early spliceosomal complexes (8). The steps involved in the execution of this novel element of the mammalian splicing code is currently not clear. Secondly, the repertoire of splicing regulatory RBPs appears to be overwhelmingly large for each of them to have the ability to interact with an early spliceosomal component. If the RBPs could promote splicing without interacting with the early spliceosomal components is not yet clear (20). Our earlier observation suggested that pre-mRNA structural remodeling by SR proteins is a possibility but the detailed mechanisms remain unclear (8). Thirdly, SR proteins may repress splicing by binding to the intronic motifs (19) through an as yet uncharacterized mechanism. Fourthly, SR proteins may bind the pre-mRNAs in multiple copies prior to assembly of the early spliceosome (6, 20). However, the mechanism that regulates whether these multiple copies of SR proteins will compete with the early spliceosomal components thus inhibiting splicing or cooperate with them for promotion of early spliceosome assembly remains unclear.

In the current work, we characterized the steps between the free pre-mRNA and the assembled early spliceosome *in vitro*. We report that different SR proteins comprising of members both having and lacking detectable interactions with the early spliceosomal components work together to promote the assembly of the early spliceosome. Our data also suggest that SR proteins and the early spliceosomal components U1 snRNP and U2AF65 have a unique competitive-cooperative relationship: binding of multiple copies of SR proteins to the pre-mRNA indeed prevent splice signal-specific recruitment of U1 snRNP and U2AF65 and instead promotes their non-specific recruitment. Interestingly, excess U1 snRNP, in the presence of U2AF65, selectively displace a fraction of the bound SR protein molecules to enable their own splice signal-specific recruitment to the pre-mRNA.

## METHODS

### Cloning, protein purification, and *in vitro* reconstitution and purification of U1 snRNP

Different subunits of U1 snRNP, SR proteins, U2AF65, U2AF35, and SF1 (1-320 a.a.) S80E/S82E were expressed and purified as described before (8). hnRNP A1 was expressed in *E. Coli* and purified as a His_6_-tagged protein, followed by further purification by gel filtration. U1 snRNP was reconstituted and purified with all full-length proteins except U1-70k, which was truncated (1-215 amino acids) (8). Additionally, a truncated form of Sm B (1-174 a.a.) was used to reconstitute U1 snRNP for certain experiments as indicated. All protein sequences used in this study are provided in the Supplementary File.

### RNA constructs

*β-globin* and *AdML* RNA were as described before (6). 14-nt long *Ron* ESE (5′-UGGCGGAGGAAGCA-3′) sequence and *AdML* 5′SS RNA (UUGGGGUGAGUACU) were described before (8, 21). *β-globin* and *AdML* RNA sequences are provided in the Supplementary File.

### Electrophoretic mobility shift assay (EMSA)

Complexes for EMSAs were assembled with the indicated concentrations of components in 20 mM HEPES-NaOH pH 7.5, 250 mM NaCl, 2 mM MgCl_2_, 1 mM DTT, 0.3% poly (vinyl alcohol) (Sigma P-8136), 0.5 M urea, and 20% glycerol. EMSA with full-length pre-mRNA and 14-nt long RNAs were carried out with 4% (89:1) and 6% (80:1) native polyacrylamide gels, respectively, as described before (8). Antibody super-shift was carried out with anti-U1-C antibody (Abcam, ab157116) and anti-SRSF1 antibody (Life Technologies, 32-4500). After the formation of complexes as described above, 0.25 µg antibody was added to the 15 µl reaction, incubated at 30 °C for 5 min and resolved on polyacrylamide gels. Bands were quantified after subtracting the background by Fiji (22). The concentration-response curves were fitted in Graphpad Prism. Our EMSAs were designed to examine both the affinity (when RNA is in trace amount) and the stoichiometry (when RNA is not in trace amount) of protein components required to form a complex following principles described before (23). Hill slope of SRSF1 binding was calculated from the percentage of bound RNA by using the equation for “log(agonist) vs. response - variable slope” on GraphPad Prism (GraphPad Software, San Diego, California, USA, graphpad.com).

### *In vitro* <u>s</u>elective 2′ <u>h</u>ydroxyl <u>a</u>cylation followed by <u>p</u>rimer <u>e</u>xtension by <u>m</u>ut<u>a</u>tional <u>p</u>rofiling (SHAPE-MaP)

For SHAPE assay with *AdML*, 500 nM refolded RNA with 3X MS2-binding loops at the 3′ end was first bound to 1500 nM MBP-MS2 protein for 10 min at 30 °C. Then the RNA was added to 250 µl reaction mixture with the same buffer composition as described before (8) at a final concentration of 100 nM, was mixed with 300 nM U1 snRNP (stock concentration 1 µM) and/or 500 nM SRSF1-RE (full-length SRSF1 with all serine residues in its RS domain replaced with glutamate) (stock concentration 8 µM), and was incubated at 30 °C for 5 min. Then complexes were bound to 15 µl amylose agarose (NEB) by rotating the tube at 4 °C for 30 min. The resin was washed with chilled buffer (20 mM HEPES pH 7.5, 250 mM NaCl, 2 mM MgCl_2_, 1 mM 2-mercaptoethanol) and then eluted in 500 µl of this buffer containing 20 mM maltose by rotating the tube at 4 °C for 2 hours. Then 250 µl solution was transferred to 8.33 µl 120 mM NMIA (final NMIA concentration 4 mM) or equal volume of DMSO (vehicle), mixed gently, and incubated at 16 °C for 1 hour. The solution was then treated with 0.5 mg/ml proteinase K in the presence of added 2 M urea at 37 °C for 30 min and the RNA was ethanol-precipitated followed by RNA cleanup with Monarch RNA cleanup kit.

SHAPE experiment shown in Figure 5C was carried out with 10 mM NMIA. The reaction mixtures contained 25 nM pre-mRNA, 125 nM or 250 nM SRSF1-RE, 600 nM U1 snRNP, 0 nM or 50 nM [U2AF65 + SF1_320_ (E/E)] (stock concentration 10 µM), and 0 or 50 nM U2AF35 (stock concentration 5 µM). Since binding of U2AF65 and U2AF35 were weak, these complexes were not purified prior to SHAPE reaction.

Preparation of denatured control with 4 mM and 10 mM NMIA, library preparation, deep sequencing, and data processing were carried out as described before (8). SHAPE-derived secondary structure models were obtained using RNAstructure software (24). All negative values in SHAPE reactivity were considered zero. For visualization of SHAPE differentials on a plot, we used fold changes in SHAPE reactivity instead of showing increase or decrease. This is because changes in SHAPE reactivity in nucleotides with inherently low SHAPE reactivity are expected to be low, but calculation of fold changes will give equal importance to the changes in SHAPE reactivity of these nucleotides as the nucleotides with inherently high SHAPE reactivity. For calculation of fold changes, all zero values were converted to 0.0001. Most fold changes are calculated by dividing the SHAPE reactivity obtained under the test condition with that of the protein-free substrate. Fold changes shown in Figure 5C are calculated by dividing the SHAPE reactivity of the complex formed in the presence of U2AF65+SF1+U2AF35 with the reactivity of the complex formed in the absence of U2AF65+SF1+U2AF35.

### Detection of RNA-protein interactions by RNP-MaP

RNP-MaP for detection of protein interaction sites on an RNA was carried out as described before (25). The early spliceosomal complexes were assembled in the presence of 0.5 M urea and U1 snRNP storage buffer contains 15 mM each arginine-HCl and glutamate-KOH. Both urea and free amino acids would interfere with the SDA crosslinking reagent used for RNP-MaP since SDA reacts with primary amines. Thus, purification of the complexes was essential for RNP-MaP analysis. Since we could purify *AdML* complexes efficiently by amylose pull-down, we carried out RNP-MaP with *AdML*. *AdML* complexes were assembled and purified as in SHAPE. The complex was eluted by rotating the tubes for 2 hours at 4 °C in elution buffer containing 4 mM SDA (NHS-diazirine, succinimidyl 4,4′-azipentanoate, Thermo Fisher Scientific, diluted from 100 mM stock in DMSO) or equivalent concentration of DMSO. Then the resin was removed by gentle centrifugation, the liquid was transferred to a 24-well plate on ice and was exposed to 3 J/cm^2^ of 365 nm light in UVP 1000CL (Analytik Jena) crosslinker. The proteins were then digested in the presence of 1.5% SDS, 20 mM EDTA, and 0.5 mg/ml proteinase K for 2 hours at 37 °C, and the RNA was purified by phenol:chloroform extraction and ethanol precipitation. The RNA pellet was dissolved in 50 µl water and was further purified by Monarch RNA cleanup kit. The reverse transcription reaction was carried out as described before (25). The library preparation and deep sequencing were carried out as in SHAPE. Analyses of RNP sites and RNP reactivities were carried out as described before (25).

### Treatment of cells with morpholino oligonucleotides, transfection-based splicing assay, and RNA immunoprecipitation

*HeLa* cells were grown in a six-well plate for transfection-based splicing assay and in 150 mm plates for RNA immunoprecipitation (RIP). Cells were grown up to 90% confluency and then transfected with 5 nmol morpholino oligonucleotide for splicing assay and 75 nmol for RIP. After 16 hours, cells were transfected with 1 µg and 15 µg *AdML* minigene plasmid, respectively. Cells were harvested 24 hours after the second transfection.

For splicing assay, cells were harvested by directly adding Trizol reagent (Thermoscientific) to the six-well plate for extraction of RNA using manufacturer’s instructions. Reverse transcription reaction was carried out with 100 ng total RNA using *AdML*-specific reverse primer in 20 µl volume, and 1 µl of cDNA preparation was used in 25 µl PCR mix.

For RIP, adherent cells were gently washed with PBS stored at room temperature, harvested with a scraper, resuspended in 30 ml PBS containing 0.1% fresh formaldehyde, and gently rocked for 10 minutes at room temperature. Then 3 ml of quenching buffer (2.5 M Glycine, 2.5 M Tris) was added to 30 ml suspension and rocked for an additional 5 min. Cells were centrifuged at 400 g for 10 min at 4 °C and washed once with chilled PBS. Cells were gently and thoroughly resuspended in chilled RIPA buffer (25 mM Tris pH 7.5, 150 mM NaCl, 1% NP-40, 0.1% SDS, 0.1% sodium deoxycholate, and 1 mM EDTA) containing 1:100 mammalian protease inhibitor cocktail (Sigma) and 1:100 RNase Inhibitor (Biobharati Life Science). Resuspended cells were briefly sonicated, and the cell lysate was cleared by centrifugation at 16000 g for 10 min at 4 °C. Cell lysate was precleared with control antibody (1 µg antibody per 100 µg of total protein) using protein-A-coated Dynabeads (Thermoscientific) following manufacturer’s instructions. Two aliquots of pre-cleared cell lysate each containing 50 µg total protein were saved for RNA extraction using the method described below and estimation of SRSF1 expression level by western blotting. Next, SRSF1 or U2AF65 was immunoprecipitated using specific antibody (1 µg antibody per 100 µg total protein) from pre-cleared cell lysate at 4 °C. Finally, the beads were separated and washed with chilled RIPA buffer.

For RNA extraction, the beads (bound to immunoprecipitant from cell lysate containing 500 µg total protein) were resuspended in 100 µl proteinase K buffer (50 mM Tris-HCl pH 7.5, 5 mM EDTA, 1% SDS, and 10 mM DTT). The pre-cleared cell lysate (50 µg total protein) was diluted in equal volume of 2X proteinase K buffer. Proteinase K was added to a final concentration of 2 mg/ml and then the solution was incubated at 65 °C for 45 min with occasional mixing to digest and de-crosslink the protein. RNA was extracted once with phenol:chloroform:isoamyl alcohol and then once with chloroform, followed by purification by Monarch RNA cleanup kit. The total RNA eluant is then digested with DNase I (NEB) for 30 min at 37 °C and purified further by Monarch RNA cleanup kit. Loss of nucleic acids before and after DNase I treatment was not detectable. 1 µl RNA without concentration adjustment was used in 20 µl reverse transcription reaction. 1 µl of cDNA preparation was used in 25 µl PCR mix.

We used αSRSF1 (rabbit IgG, A302-052A, Bethyl Laboratories) (26) and control antibody (Rabbit IgG, Diagenode c15410206) for immunoprecipitation of SRSF1, and αU2AF65 (mouse IgG2b, Sigma, U4758) (3) and control antibody (mouse IgG2b, Cell Signaling, 53484) for immunoprecipitation of U2AF65.

### Amylose pull-down assay

Pre-mRNA with 3X MS2-binding loop at its 3′ end was mixed with 1X molar excess MBP-MS2 (to keep the MBP-MS2 band intensity low so that it does not mask other bands with similar migration patterns) and incubated for 10 min at 30 °C. Then this RNA was mixed with the indicated components in 100 µl volume. The buffer composition for the assembly reaction was same as described earlier (8). After each addition, the reaction was incubated at 30 °C for 5 min. Then the reaction was incubated at room temperature for 5 min and then placed on ice for 5 min. Next, it was centrifuged at 16000 g for 15 min at 4 °C. The supernatant was then transferred to a fresh low-bind tube containing pre-washed 5 µl amylose magnetic beads (NEB) and rotated for 45 min at 4 °C. Magnetic beads, due to highly smooth surfaces, suppress non-specific beads:protein interactions. Then the supernatant was removed, and the beads were washed thrice with 400 µl chilled wash buffer (20 mM HEPES-NaOH pH 7.5, 250 mM NaCl, 1 mM DTT, 2 mM MgCl_2_, and 1 M urea). The beads were then boiled in Laemmli buffer and analyzed on 15% SDS gel. For RNA analysis, the beads were resuspended in proteinase K buffer (50 mM Tris-HCl pH 7.5, 5 mM EDTA, 1% SDS, and 10 mM DTT) in the presence of 0.5 mg/ml proteinase K and was incubated at 37 °C for 30 min. After removal of the beads, RNA was purified by phenol:chloroform:isoamyl alcohol extraction and ethanol precipitation. Then the RNA pellet was resuspended in deionized formamide, heated for 2 min at 95 °C, resolved by urea PAGE, and stained with 0.1% toluidine blue. Densitometric quantification of band intensity was carried out after background subtraction using Fiji (22).

## RESULTS

### U1 snRNP recognizes the pre-mRNA 3D structural scaffold in *β-globin* pre-mRNA

Our earlier results revealed that early spliceosomal components recognize the 3D structural scaffold of adenovirus 2 major late transcript IVS I (*AdML*) (8). Here we expanded on the observation and examined if similar mechanisms are employed for the recognition of human *β-globin* IVS1.

We titrated 10 pM *β-globin* in the absence or presence of 60 nM SRSF1-RBD (Supplementary Figure S1A) with U1 snRNP (Supplementary Figure S1B, Figure 1A). Without SRSF1-RBD, U1 snRNP formed a single complex with *β-globin* (lanes 7-8). In the presence of SRSF1-RBD, U1 snRNP formed multiple complexes. We confirmed the presence of SRSF1-RBD as well as U1 snRNP in the final WT ‘ternary’ complexes by performing antibody super-shift with anti-SRSF1 and anti-U1-C antibodies, respectively (Figure 1A – compare lanes 5 & 9 and 18 & 19). Anti-U1C antibody disintegrated a portion of the pre-mRNA complex likely because U1-C is important for RNA binding of U1 snRNP (27). Band intensity of the free probes suggests the efficiency of complex formation: a lower level of probe remains free in the presence of SRSF1-RBD than in its absence, which suggests that SRSF1-RBD stabilizes U1 snRNP binding to *β-globin* (Figure 1A, compare lanes 4 & 5 with 7 & 8).

**Figure 1.**
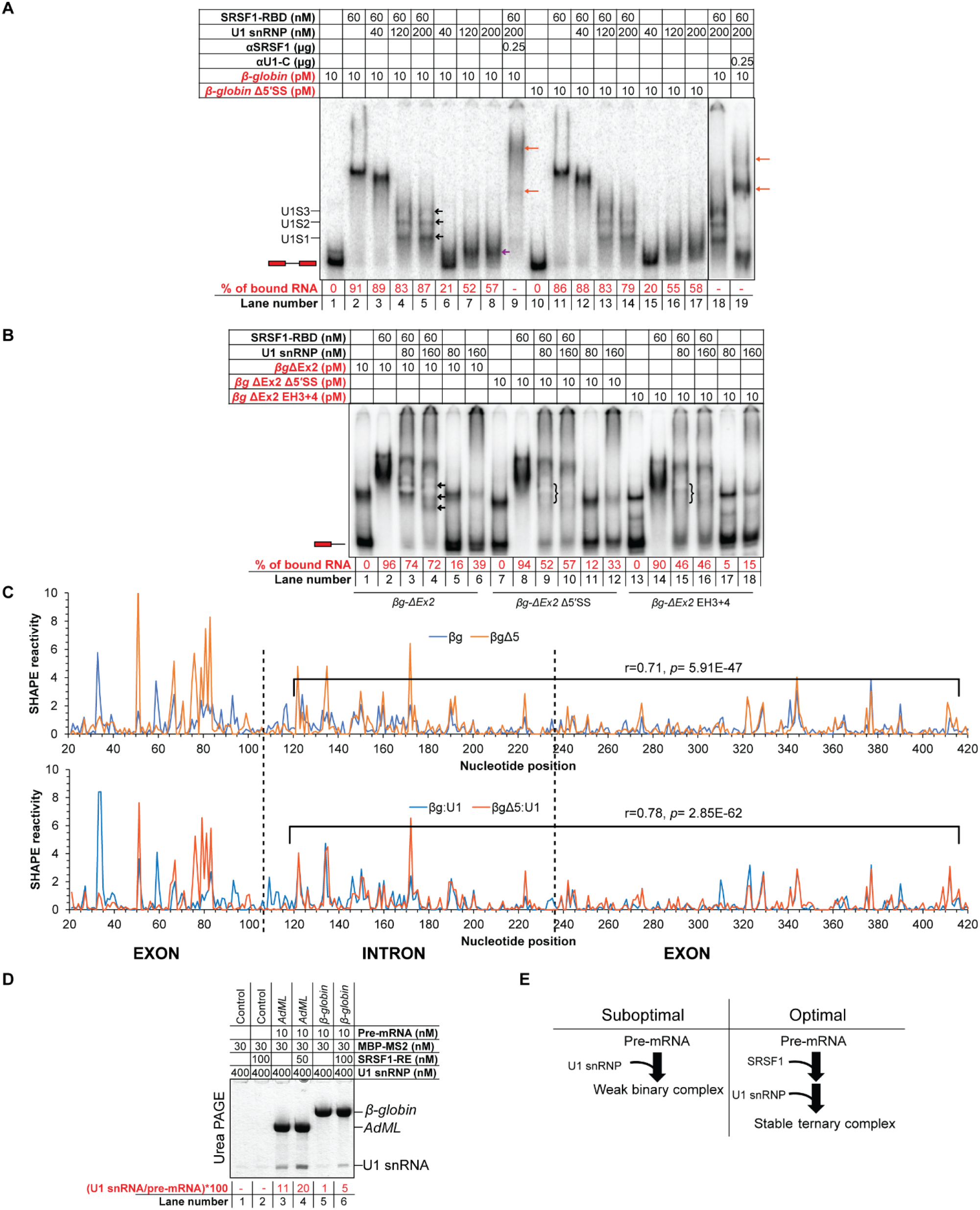
U1 snRNP recognizes the pre-mRNA 3D structural scaffold in *β-globin*. (A) *β-globin* forms ternary complexes with SRSF1-RBD and U1 snRNP, which migrate primarily as three major bands (marked with black arrows in lane 5 and labeled as U1S1, U1S2, and U1S3 on the side of the gel); U1 snRNP also binds free *β-globin* (marked with a violet arrow in lane 8); the ternary complex is super-shifted with αSRSF1 (lane 9, marked with orange arrows) and αU1-C (lanes 18-19); slightly more smeary complexes of similar migration patterns were formed with *β-globin* Δ5′SS (lanes 10-17); red script indicates radiolabeled components; the position of the free probe is indicated with an exon-intron-exon schematic; percentage of upshifted *β-globin* probe is indicated below each lane. (B) *βg-ΔEx2* (*β-globin* lacking the 3′ exon) formed U1 snRNP-dependent complexes in the presence of 60 nM SRSF1-RBD (the complexes are marked with arrows in lane 4); these U1 snRNP-dependent complexes were significantly weakened for *βg-ΔEx2* with 5′SS mutations (*βg-ΔEx2* Δ5′SS) and *βg-ΔEx2* with hybridization-mutation immediately upstream of the 5′SS (*βg-ΔEx2* EH3+4) (the position of the weakened complexes are marked with curved brackets in lanes 9 & 15); U1 snRNP shows no well-defined interaction with the free RNAs (lanes 5, 6, 11, 12, 17, 18); the position of the free probe is indicated with an exon-intron schematic. (C) Overlaid SHAPE reactivity of protein-free *β-globin* WT and its 5′SS mutant (Δ5) (top) and U1 snRNP-bound *β-globin* WT and U1 snRNP-bound *β-globin* 5′SS-mutant (bottom); the segments showing a moderate correlation of SHAPE reactivity are marked in each plot and the corresponding r and *p* values are indicated; exon-intron boundaries are demarcated with dotted vertical lines. (D) Amylose pull-down assay showing enhancement of co-purification of U1 snRNP (indicated by corresponding U1 snRNA level) with *AdML* and *β-globin* in the presence of SRSF1-RE by urea PAGE. (E) Summary flow chart: U1 snRNP specifically recognizes a global 3D structural scaffold of *β-globin* and SRSF1 enhances this interaction.

*β-globin* Δ5′SS, which has 14 substitution mutations in the authentic 5′SS and two cryptic sites at the −16 and +13 positions (28) formed similar complexes with U1 snRNP and/or SRSF1-RBD as the WT substrate (Figure 1A - compare lanes 4, 5 with 13, 14 and lanes 7, 8 with 16, 17, respectively). We hypothesized that the 5′SS sequence plays a relatively minor role in stabilizing the early complex and that the long 3′ exon of *β-globin* (206-nt in *β-globin*) might provide sufficient support to stably recruit U1 snRNP. Therefore, we generated a *β-globin* construct with a truncated 3′ exon (*βg-ΔEx2*) and introduced Δ5′SS mutations into this construct to generate *βg-ΔEx2* Δ5′SS. We also generated the hybridization-mutant (EH3+4) on the *βg-ΔEx2* construct, in which the critical single-stranded segments immediately upstream of the 5′SS (6) are hybridized; this mutant does not recruit SR proteins appropriately and is defective in splicing (6). These probes migrate as two bands – the majority as the fast-migrating natively folded RNA and the minority as the slow migrating unfolded/misfolded RNA (29) (Figure 1B – lane 1). U1 snRNP alone did not form a stable complex with *βg-ΔEx2* (lanes 5, 6). However, in the presence of SRSF1-RBD, U1 snRNP formed the distinctively migrating complexes (lanes 3, 4). The ternary complex assembled with *βg-ΔEx2* is less stable than the same with the full-length probe (*β-globin*) as suggested by the remaining free probe level (compare lanes 4, 5 of Figure 1A with 3 & 4 of Figure 1B). Nevertheless, the ternary complexes formed with *βg-ΔEx2* were more stable than those formed with the other two mutant probes (compare lanes 3 & 4 with 9 & 10 and 15 & 16). These results strongly suggest that the entire length of the pre-mRNA including the 5’SS is essential for U1 snRNP recruitment and that SRSF1 stabilizes the complex.

The fact that the 3′ exon stabilizes U1 snRNP on *β-globin* suggests that U1 snRNP recognizes a 3D structural scaffold that integrates the 3′ exon of *β-globin*. To examine how the *β-globin* structure regulates U1 snRNP binding even in the absence of 5′SS, we examined the SHAPE reactivity of protein-free *β-globin* and its 5′SS-mutant and generated their SHAPE-derived secondary structure models (Supplementary Figure S1C, S1D). The largest structural differences were observed around the 5′SS between the 50^th^ to 120^th^ positions (5′ exon ends at the 106^th^ position) (Figure 1C – top panel). However, the strandedness of nucleotides in the remainder of the pre-mRNA appeared somewhat similar. SHAPE reactivity of nucleotides between the 120^th^ and 420^th^ positions of both RNAs showed a moderate correlation (0.71). SHAPE reactivity of U1 snRNP-bound *β-globin* variants exhibited a slightly higher correlation between the 120^th^ and 420^th^ positions (0.78) (Figure 1C – bottom panel). This suggests that mutation in the authentic and cryptic 5′SS did not completely disrupt the structure of *β-globin* and that a global interaction of U1 snRNP with *β-globin* is important for U1 snRNP recruitment to this splicing substrate. Additionally, we observed a loss of SHAPE reactivity over 1000-fold at several positions in the 5′SS region and its flanking segments in both the mutant and the WT substrate upon binding of U1 snRNP; among these nucleotides, the 104^th^ nucleotide at the −3 position of the 5′SS, which is not mutated in Δ5′SS, and the 58^th^ and the 47^th^ nucleotides (also not mutated) within the critical single-stranded segment immediately upstream of the 5′SS, are common to both substrates (Supplementary Figure S1E). This likely suggests that the 3D structural scaffold of *β-globin* Δ5′SS is able to position U1 snRNP in a similar manner as the WT substrate. However, the SHAPE reactivity of *β-globin* and *β-globin* + U1 snRNP exhibits a lower correlation (0.77; *p* = 3.12E-60) compared to that of *β-globin* Δ5′SS and *β-globin* Δ5′SS + U1 snRNP (0.87; *p* = 2.57E-93). This suggests that U1 snRNP contacts and the consequent structural remodeling of the WT substrate is more extensive than the mutant substrate. Nonetheless, as expected, *β-globin* Δ5′SS is completely defective in splicing *in vivo* (Supplementary Figure S1F).

The results shown so far suggest that SRSF1 stabilizes the U1 snRNP: *β-globin* complex. To further confirm this observation, we performed an amylose pull-down assay with 3XMS2-tagged *β-globin* in the presence of SRSF1-RE and U1 snRNP. We also used 3XMS2-tagged *AdML*, which binds U1 snRNP more strongly (8). From the pulled-down samples, we isolated the total RNA and analyzed the proportion of pre-mRNA and U1 snRNA in each sample. This revealed co-purification of more U1 snRNA in the presence of SRSF1-RE with both substrates (Figure 1D).

Overall, these results suggest that U1 snRNP specifically recognizes a global 3D structural scaffold of *β-globin* and that SRSF1 enhances this interaction (Figure 1E). This scaffold exhibits a nominal alteration in its structure and in its ability to be recognized by U1 snRNP upon mutation of the authentic and cryptic 5′SS.

### Cooperative binding of SRSF1 in appropriate stoichiometry to the pre-mRNA is important for stable recruitment of U1 snRNP

To understand how SRSF1 enhances the U1 snRNP:pre-mRNA interactions as shown above and reported previously, we characterized the nature of binding of SRSF1 to the pre-mRNA. Upon titration of 10 pM radiolabeled *β-globin* with SRSF1-RBD, we observed that the most compact band on native gel was formed with ∼ 40 nM protein, and at higher protein concentrations, larger, slower migrating complexes were formed (Figure 2A). The generation of a concentration-response curve with densitometrically-quantified free probe intensity showed the Hill coefficient to be much greater than 1.0, suggesting that SRSF1-RBD binding is cooperative (Figure 2B). In contrast to this high-affinity and cooperative binding, K_d_ of the SRSF1-RBD-ESE complex is around 0.5 µM (21). Consistently, we also found that SRSF1-RBD weakly binds to a 14-mer probe containing the *Ron* ESE site, a strong binding site for SRSF1 (21) (Figure 2C). A stoichiometric binding assay revealed that about 28 molecules of SRSF1-RBD were required to form the most compact band on the native gel (Supplementary Figure S2A). A similar result was also obtained with *AdML* pre-mRNA (Supplementary Figure S2B) with a Hill coefficient of ∼ 3 (Supplementary Figure S2C), which binds about 10 molecules of SRSF1-RBD for formation of the most compact band on the native gel (6).

**Figure 2.**
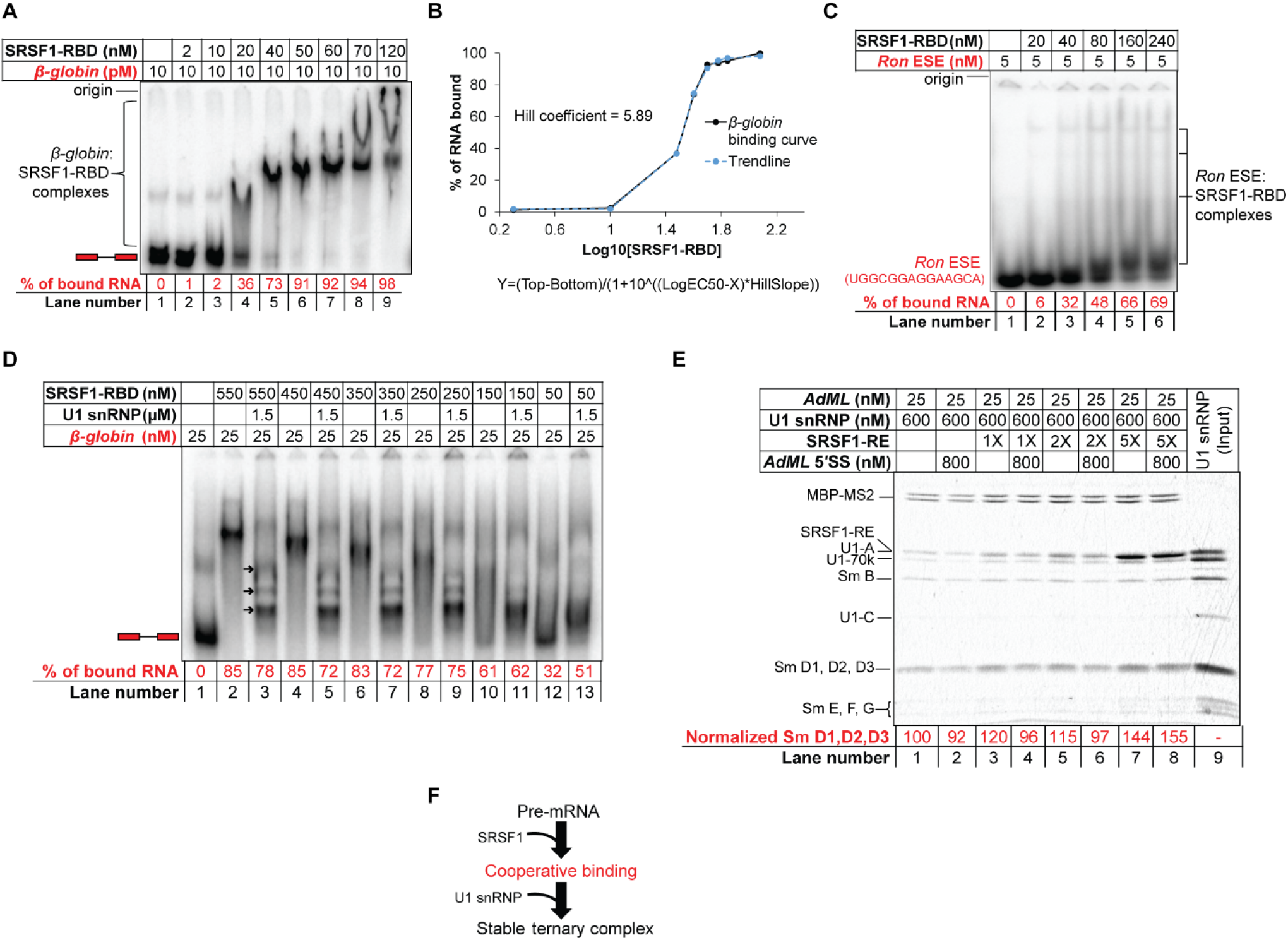
Cooperative binding of SRSF1-RBD to *β-globin* is important for stabilization of U1 snRNP. (A) EMSA showing gradual upshift of radiolabeled *β-globin* complexes upon titration with SRSF1-RBD. (B) Binding curve (solid black line) of SRSF1-RBD to *β-globin* and Hill coefficient obtained from band intensity of the free probe in A; the equation of the curve is given below it, where Y is band intensity, X is Log10[SRSF1], and Top and Bottom are plateaus in the unit of the Y axis; a trendline (broken blue line) was generated to visually examine the goodness of fit by estimating the Y values from the X-values, Hill Slope (5.89), EC50 (33), Top (98.18), and Bottom (1.85). (C) EMSA showing formation of weak complexes with a radiolabeled 14-nt long RNA containing *Ron* ESE (a binding site for SRSF1). (D) Binding efficiency of SRSF1-RBD diminishes with decreasing SRSF1:*β-globin* ratio below 10:1 (compare lanes 8 and 10) with a corresponding decline in the ternary complex formation with U1 snRNP (compare lanes 9 and 11). (E) Amylose pull-down assay of *AdML* (25 nM) bound to MBP-MS2 showing enhanced stability of U1 snRNP with 5X molar excess SRSF1-RE (125 nM) compared to 2X, 1X, or no SRSF1-RE, particularly when challenged with a 14-nt long RNA containing *AdML* 5′SS; Sm D1, D2, D3 band intensity is normalized to MBP-MS2 level. (F) Summary flow chart: Specific U1 snRNP recruitment requires cooperative binding of SRSF1, which in turn requires a threshold level of SRSF1:pre-mRNA molar ratio; red text indicates the step added based on the conclusions of this Figure.

Although *β-globin* and *AdML* require about 28 and 10 molecules of SRSF1-RBD respectively to form the most compact band visible on the native gel, we hypothesize that a lower SRSF1:pre-mRNA ratio could be sufficient to form the ternary complex with U1 snRNP efficiently. In a stoichiometric EMSA, we found that 60X molar excess U1 snRNP was required for forming the well-resolved ternary complexes with 1X *β-globin* pre-mRNA (Supplementary Figure S2D). We next performed another stoichiometric EMSA to estimate the minimum SRSF1:pre-mRNA ratio required to form stable ternary complexes with U1 snRNP. For this, we added 60X molar excess U1 snRNP to 1X *β-globin* probe bound to 2X, 6X, 10X, 14X, 18X, or 22X molar excess SRSF1-RBD molecules (Figure 2D). Quantification of the unbound *β-globin* probe in each lane suggests that an SRSF1-RBD:*β-globin* ratio of at least 10:1 is required to upshift the majority of the free probe (compare lanes 8 and 10) and form the ternary complex efficiently with U1 snRNP (lanes 9 and 11). We also examined the minimum SRSF1:*AdML* ratio required for formation of the ternary complex by amylose pull-down assay (Figure 2E). The level of bound U1 snRNP increased with an increasing SRSF1-RE:*AdML* ratio (lanes 1, 3, 5, and 7). Challenging each of these complexes with a 14-nt long RNA containing *AdML* 5′SS, which disrupts the binary *AdML*+U1 snRNP complex more efficiently than the ternary complex (8), caused a reduction in the U1 snRNP level bound to *AdML* in the presence of 0X, 1X, and 2X SRSF1-RE but not 5X SRSF1-RE (lanes 2, 4, 6, and 8). This suggests that complexes formed in the presence of 1X and 2X SRSF1-RE (lanes 2, 4, and 6) represent a mixture of binary and ternary complexes while the ones formed with 5X SRSF1-RE (lane 8) are primarily ternary complexes.

Taken together, these results suggest that SRSF1 engages the pre-mRNA in a cooperative mode, which is required for stabilization of the interactions between the pre-mRNA and U1 snRNP (Figure 2F).

### U1 snRNP selectively displaces some SRSF1 molecules bound to the pre-mRNA

To further understand how pre-mRNA, SRSF1, and U1 snRNP assemble to form the stable ternary complexes, we titrated 10 pM *β-globin* bound to about 25 molecules of SRSF1-RBD with U1 snRNP. Increasing concentrations of U1 snRNP gradually increased the mobility of the pre-mRNA complex, eventually forming the faster migrating complexes (marked with arrows in Figure 3A). We also observed that the migration rate of the αSRSF1-super-shifted complexes increased with increasing concentrations of U1 snRNP up to 80 nM (lanes 3, 5, and 7 of Figure 3B). This suggests that SRSF1-RBD is gradually depleted from *β-globin* pre-mRNA with increasing concentrations of U1 snRNP. To further confirm the displacement of SRSF1 from the full-length pre-mRNA by U1 snRNP, we performed amylose pull-down assays with MS2-tagged *β-globin* pre-mRNA substrate in a complex with SRSF1-RE or SRSF1-RE and U1 snRNP (Figure 3C). Densitometric analysis suggested that > ∼60% of bound SRSF1-RE, which stains 5X more intensely than U1 snRNP proteins (8), is displaced by U1 snRNP. We also repeated the experiment with 25 nM (1X) *AdML* WT or ΔPPT bound to 250 nM SRSF1-RBD (10X), which was titrated with 200 nM (8X) and 600 nM (24X) U1 snRNP. Densitometric analysis indicated that 8X and 24X U1 snRNP displaces ∼ 53% and ∼ 62% of SRSF1-RBD bound to *AdML* WT and ∼ 49% and ∼ 69% of SRSF1-RBD bound to *AdML* ΔPPT (Figure 3D, values normalized with MBP-MS2 levels). Interestingly, with 8X U1 snRNP, *AdML* ΔPPT bound more U1 snRNP than *AdML* WT. However, with 24X U1 snRNP, *AdML* WT bound more U1 snRNP. Given that *AdML* ΔPPT is unable to bind U1 snRNP as strongly as the WT substrate in the protein-free state due to a disrupted pre-mRNA scaffold (8), this result suggests that the displacement of SRSF1 from SRSF1:*AdML* binary complex improves the splice signal specificity of U1 snRNP recruitment (Figure 3D). We also observed that U1 snRNP displaces SRSF1-RBD from 14-nt long RNA containing *Ron* ESE, a strong binding site for SRSF1 (21) (Supplementary Figure S3A).

**Figure 3.**
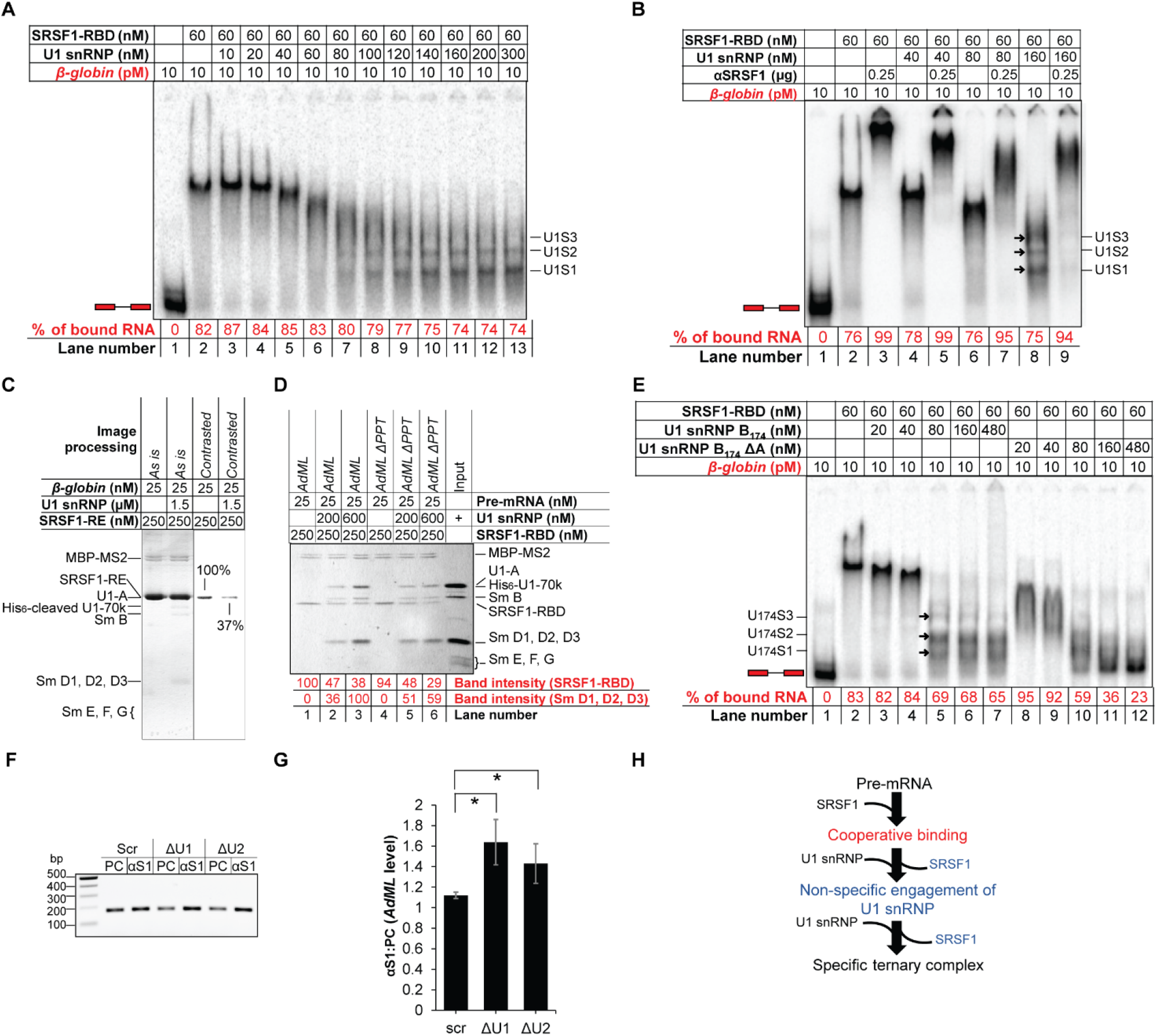
Excess U1 snRNP displaces majority of SRSF1-RBD molecules bound to the pre-mRNAs. (A) Titration of *β-globin* bound to SRSF1-RBD with U1 snRNP showing change in the migration of the radioactive complexes leading to formation of the ternary complexes (labeled on the side of the gel). (B) Super-shift of complexes shown in A with αSRSF1 showing increase in the migration rate of the super-shifted complexes up to 80 nM U1 snRNP. (C) Amylose pull-down assay of MS2-tagged *β-globin* bound to MBP-MS2 and SRSF1-RE in the presence or absence of U1 snRNP showing U1 snRNP-mediated displacement of SRSF1-RE from *β-globin*. (D) Amylose pull-down assay of MS2-tagged *AdML* WT and ΔPPT showing that U1 snRNP displaces SRSF1-RBD bound to the pre-mRNA, which improves specific interactions of U1 snRNP with the WT pre-mRNA complex more significantly than *AdML* ΔPPT. (E) U1 snRNP assembled with Sm B_1-174_ and without U1-A displaces SRSF1-RBD molecules from *β-globin* releasing free pre-mRNA (lane 12); the ternary complexes formed with U1 snRNP B_174_ are marked with arrows and labeled as U_174_S1, U_174_S2, and U_174_S3. (F) Quantification of *AdML* pre-mRNA by RT PCR using a reverse primer nested within the intron in pre-cleared extract (50 µg total protein, PC) or immunoprecipitant obtained from precleared extracts (500 µg total protein) with anti-SRSF1 antibody (αS1) from *HeLa* cells treated with scrambled (scr), U1 (ΔU1), and U2 (ΔU2) AMO. (G) Plot of the ratio of immunoprecitated *AdML* pre-mRNA and *AdML* pre-mRNA in the pre-cleared extract showing significant enrichment of *AdML* upon disruption of U1 snRNP as well as U2 snRNP; error bars indicate standard deviation, ‘*’ *p*<0.05, n=3. (H) Summary flow chart: U1 snRNP engages the SRSF1-pre-mRNA complex non-specifically and displacement of some of the SRSF1 molecules bound to the pre-mRNA by a molar excess of U1 snRNP is required to establish a specific interaction with U1 snRNP; blue text indicates the steps added based on the conclusions of this Figure.

To further demonstrate whether U1 snRNP displaces SRSF1-RBD from the full-length pre-mRNA, we sought to identify a U1 snRNP variant that can displace SRSF1-RBD from the pre-mRNA without itself getting recruited. Since different protein components of U1 snRNP play important roles in binding of U1 snRNP to the RNA (27), we assembled U1 snRNP with varied components to identify a variant that would not bind the pre-mRNA. We found that assembling U1 snRNP with Sm B_1-174_ and without U1-A (U1 snRNP B_174_ ΔA) served this purpose. We then titrated SRSF1-RBD-bound *β-globin* with U1 snRNP B_174_ and U1 snRNP B_174_ ΔA (Figure 3E). While U1 snRNP B_174_ formed complexes of comparable migration patterns as observed with U1 snRNP (lanes 5, 6, 7), U1 snRNP B_174_ ΔA at high concentrations displaced all SRSF1-RBD molecules from *β-globin* releasing the free probe (lane 12). To examine if U1 snRNP B_174_ ΔA displaces SRSF1-RBD from the pre-mRNA due to a higher affinity for SRSF1-RBD, we tested the assembly of the binary U1 snRNP:SRSF1-RBD complex by anion-exchange chromatography (Supplementary Figure S3B). While U1 snRNP B_174_ formed a stoichiometrically 1:1 binary complex that eluted under a sharp and symmetric peak (Supplementary Figure S3B – top, S3C – top), U1 snRNP B_174_ ΔA did not (Supplementary Figure S3B – bottom, S3C – bottom). We found that U1 snRNP B_174_ ΔA also displaces SRSF2-RE and SRSF5-RBD from *β-globin*, both of which promote its splicing (30) (Supplementary Figure S3D)

To examine if SRSF1 displacement is in the early spliceosome assembly pathway in cells, we inhibited assembly of U1 snRNP and U2 snRNP by transfecting *HeLa* cells with a 25-nt long morpholino oligonucleotide that is complementary to the 5′ end of U1 snRNA (U1 AMO) or U2 AMO since destabilization of U2 snRNP also inhibits stable recruitment of U1 snRNP without directly impairing the integrity of U1 snRNP (31). Cells were then transfected with *AdML* minigene and its splicing efficiency was tested by RT PCR: U1 and U2 AMO partially inhibited splicing compared to the scrambled AMO (Supplementary Figure S3E). SRSF1 was then immunoprecipitated from the pre-cleared cell lysate and *AdML* pre-mRNA levels in the immunoprecipitant were quantified by RT PCR (Figure 3F and 3G, Supplementary Figure S3F). The level of enrichment of *AdML* pre-mRNA in the immunoprecipitant compared to the precleared cell lysate was highest for U1 AMO and lowest for the scrambled AMO suggesting that disruption of U1 snRNP and U2 snRNP leads to a more persistent interaction between SRSF1 and *AdML* pre-mRNA. To examine the level of SRSF1 in the precleared cellular extract, which would dictate the level of immunoprecipitated SRSF1 and the RNA associated with it, we examined SRSF1 expression levels in precleared extracts. SRSF1 band intensity normalized with the internal control GAPDH suggested that cells express similar levels of SRSF1 upon any of the treatments (Supplementary Figure S3G, H, I).

Overall, these results suggest that U1 snRNP selectively displaces some SRSF1 molecules bound to the full-length pre-mRNA prior to its own specific recruitment to the pre-mRNA (Figure 3H).

### RNP reactivity verifies SR protein-mediated pre-mRNA structural remodeling for the stable recruitment of U1 snRNP

We next investigated the sites of ‘optimal’ binding of SRSF1 and its effects on pre-mRNA structure and U1 snRNP recruitment *in vitro*. Purification of the pre-mRNA:SR:U1 snRNP complexes was required for the SHAPE and RNP-MaP analyses (see Methods), which led us to use *AdML* pre-mRNA. We compared fold changes in SHAPE reactivity of *AdML* upon binding of 5X SRSF1-RE (SHAPE_5XS1, the optimal SRSF1 stoichiometry), 2X SRSF1-RE (SHAPE_2XS1 – suboptimal SRSF1 stoichiometry), U1 snRNP (SHAPE_U1), and 5X SRSF1-RE + U1 snRNP (SHAPE_5XS1+U1). SHAPE_5XS1 exhibits a greater enhancement of nucleotide flexibility across the pre-mRNA compared to SHAPE_2XS1 (Figure 4A). Enhanced flexibility of segments immediately upstream of the 5′SS (marked with a rectangle above the plot) and at positions 86 and 129 (marked with arrows above the plot) is lessened in SHAPE_5XS1+U1 (Figure 4B). This suggests that SRSF1-mediated remodeling enhances flexibility of these segments to promote the likely contacts with U1 snRNP. Of these segments, the one immediately upstream of the 5′SS is particularly important for splicing across the transcriptome (6). Finally, we compared the fold changes in SHAPE reactivity in SHAPE_U1 and SHAPE_5XS1+U1 (Figure 4C). SHAPE_5XS1+U1 exhibited a much higher level of flexibility-gain across the pre-mRNA compared to SHAPE_U1. These observations are consistent with our previous report that splicing-competent pre-mRNAs are highly flexible *in vivo* (6) correlating the SR protein-mediated pre-mRNA structural remodeling observed *in vitro* with the *in vivo* structural state of the pre-mRNA.

**Figure 4.**
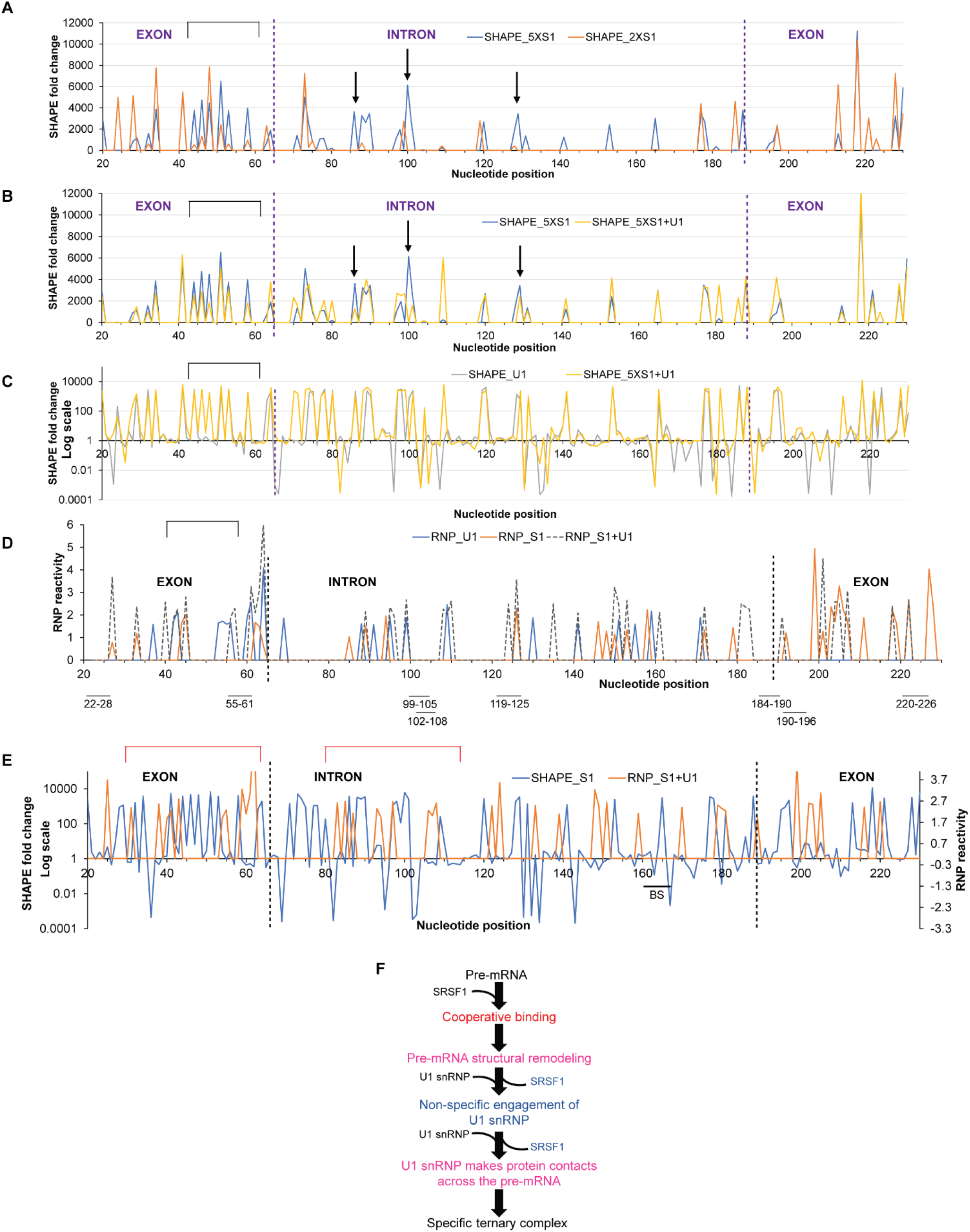
Global structural remodeling and protein contacts involved in U1 snRNP recruitment to *AdML*. (A) Fold changes in SHAPE reactivity of *AdML* upon binding of 5X SRSF1-RE (optimal stoichiometry, blue line) or 2X SRSF1-RE (sub-optimal stoichiometry, orange line); fold changes in SHAPE reactivity are plotted along *y*-axis showing only the increase in SHAPE reactivity (i.e., *y* >1); the nucleotides that exhibit a gain of flexibility over 1000 fold with 5X SRSF1-RE but not 2X SRSF1-RE and a loss of flexibility with [5X SRSF1-RE + U1 snRNP] shown in B are marked with either a rectangle or arrows above the plot. (B) Fold changes in SHAPE reactivity upon binding of 5X SRSF1-RE (blue line) or [5X SRSF1-RE + U1 snRNP] (yellow line); fold-changes in SHAPE reactivity are plotted along *y*-axis showing only the increase (i.e., *y* > 1). (C) Fold changes in SHAPE reactivity upon binding of U1 snRNP alone (grey line) or [5X SRSF1-RE + U1 snRNP] (yellow line); *y*-axis is shown in Log_10_ scale. (D) RNP reactivity of *AdML* bound to U1 snRNP (blue line), 5X SRSF1-RE (orange line), and [5X SRSF1-RE + U1 snRNP] (grey dotted line); *in vitro* selected SRSF1-binding sites are demarcated below the plot and exon-intron boundaries are indicated with dotted vertical lines. (E) Fold changes in SHAPE reactivity of [5X SRSF1-RE + *AdML*] (blue line) plotted along primary *y*-axis and RNP-reactivity of [5XSRSF1 + U1 snRNP + *AdML*] (orange line) along secondary *y*-axis. (F) Summary flow chart: Cooperative binding of SRSF1 followed by displacement of some of the SRSF1 molecules by excess U1 snRNP exposes an optimally remodeled pre-mRNA to specifically engage U1 snRNP through multiple contacts across the pre-mRNA scaffold; pink text indicates the steps added based on the conclusions of this Figure.

We then examined the protein contacts with the RNA by RNP-MaP-seq, which identifies the nucleotides within ∼ 12 Å of the central carbon atom of an interacting lysine residue, under three different binding conditions: 5XSRSF1 alone (RNP_S1), U1 snRNP alone (RNP_U1), and 5X SRSF1 and U1 snRNP (RNP_S1+U1) (Figure 4D). The protein contacts were also mapped onto the SHAPE-derived secondary structure model of *AdML* pre-mRNA (Supplementary Figure S4A, B, C). The following observations were made from the RNP experiments: (1) RNP_U1 exhibits protein contacts across exon 1 and the intron only (23 sites), while RNP_S1 and RNP_S1+U1 across the entire pre-mRNA (44 and 40 sites, respectively). (2) 25 RNP-reactive nucleotides in RNP_S1 were not reactive in RNP_S1+U1 likely due to the displacement of SRSF1 from the pre-mRNA. (3) 17 out of 19 common RNP-reactive sites in RNP_S1 and RNP_S1+U1 exhibit a stronger interaction (i.e., higher RNP reactivity) in RNP_S1+U1. (4) Similarly, 9 out of 10 RNP-reactive nucleotides common to both RNP_U1 and RNP_S1+U1 exhibit a stronger interaction in RNP_S1+U1. (5) Three nucleotides (41^st^, 62^nd^, and 124^th^) exhibit RNP reactivity in all three samples and the reactivity is the highest in RNP_S1+U1. (6) Upon comparison of SHAPE_S1 and RNP_S1+U1, we observed that segments flanking the 5′SS that have enhanced flexibility in SHAPE_S1 exhibit multiple lysine contacts in RNP_S1+U1 (Figure 4E).

Overall, these results suggest that both binding and displacement of SRSF1 to the pre-mRNA play important roles in structural remodeling of the pre-mRNA regulating global engagement of U1 snRNP (Figure 4F).

### A precise SRSF1:pre-mRNA ratio is required for specific recruitment of U2AF65

To examine the effects of the SRSF1 level bound to the pre-mRNA on U2AF65 recruitment, we performed amylose pull-down assays with 25 nM MS2-tagged WT *AdML* pre-mRNA and its PPT mutant bound to 1X, 2X, and 5X molar excess SRSF1-RE and reacted with 24X molar excess U1 snRNP and 50 nM [U2AF65 + SF1_320_ (E/E)] (Figure 5A, Supplementary Figure S5A). U2AF65 binding increased with an increasing SRSF1:pre-mRNA ratio for *AdML* WT but remained low for *AdML* ΔPPT at a so-called background level. This suggests that a minimum SRSF1:pre-mRNA ratio is critical for U2AF65 recruitment to *AdML* as this ratio dictates cooperative binding of SRSF1 to the pre-mRNA. Additionally, in the presence of U2AF65, an increase in the bound U1 snRNP level was not observed with the increasing SRSF1:pre-mRNA ratio (Figure 5A) unlike in the absence of U2AF65 (Figure 2E). To examine the reason for this, we performed amylose pull-down assay of *AdML* mixed with 5X SRSF1-RBD and 24X U1 snRNP in the presence as well as the absence of 50 nM [U2AF65 + SF1_320_ (E/E)] (Supplementary Figure S5B). We used SRSF1-RBD instead of SRSF1-RE because it is possible to quantify SRSF1-RBD band intensity accurately. We found lesser levels of SRSF1-RBD and U1 snRNP bound to *AdML* in the presence of U2AF65. Thus, the displacement of SRSF1 from the pre-mRNA, a process that diminishes non-specific interactions of U1 snRNP with the pre-mRNA (Figure 3D), is more efficient when both U1 snRNP and U2AF65 are present. Nonetheless, we did not observe binding of U2AF65 to the pre-mRNA in the presence of SRSF1-RBD (Supplementary Figure S5B). This suggests that stabilization of U2AF65 by SRSF1-RBD is not sufficiently strong to be detected by the pull-down assay but by the EMSA as shown later (Figure 5E) and that possibly the phosphorylated RS domain of SRSF1 and the RS domain of U2AF65 interact for engaging the branchsite (32).

**Figure 5.**
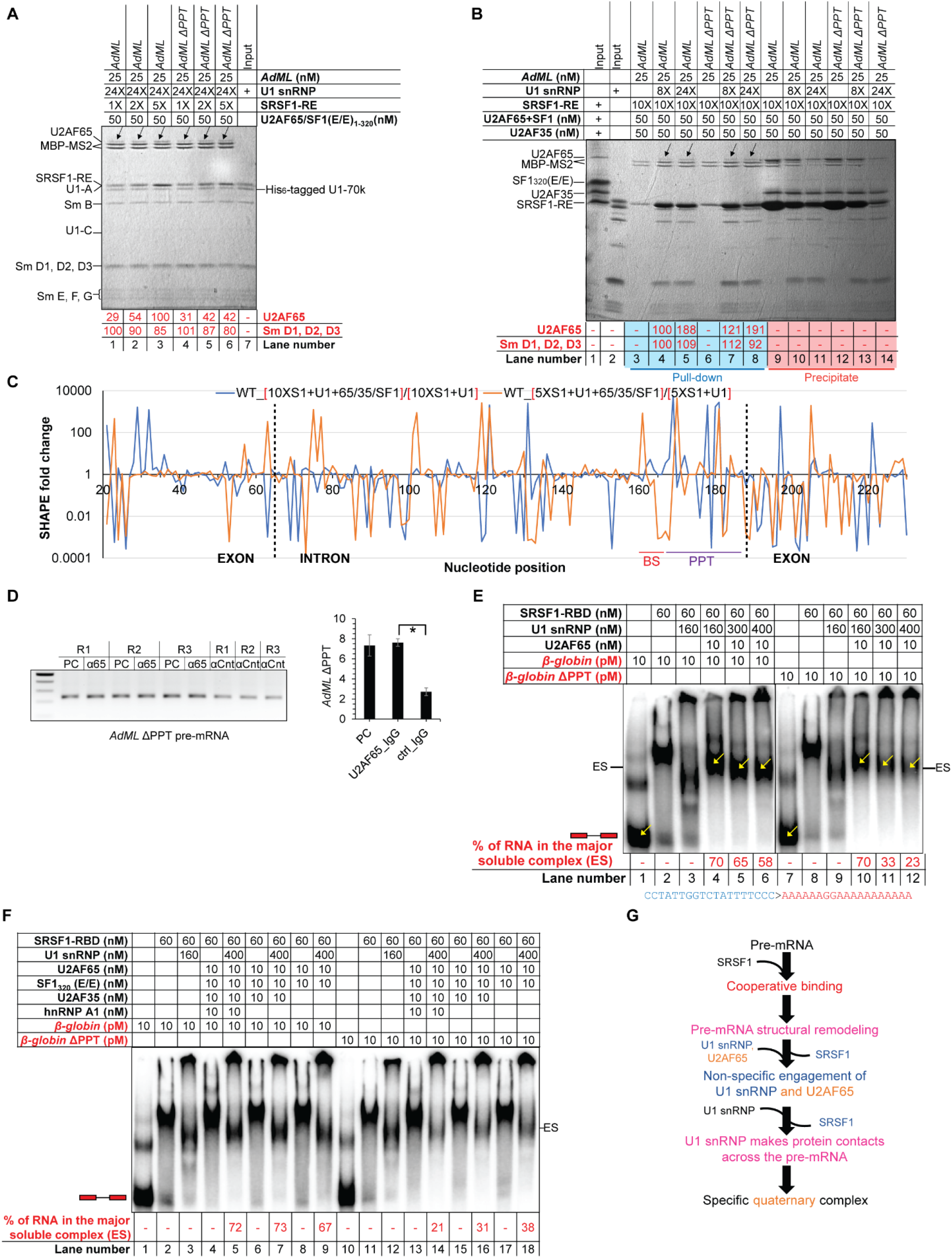
Specific U2AF65 recruitment requires cooperative binding and displacement of SRSF1. (A) Amylose pull-down assay showing binding of increasing levels of U2AF65 with increasing SRSF1-RE:*AdML* ratio but fixed U1 snRNP:*AdML* ratio; black arrows mark the U2AF65 bands; band intensities normalized to the MBP-MS2 levels are shown; a background-subtracted and contrasted version of this image is shown in Supplementary Figure S5A. (B) Amylose pull-down of *AdML* in the presence of 8X or 24X molar excess U1 snRNP, 10X molar excess of SRSF1-RE, and 50 nM [U2AF65 + SF1_320_ (E/E) + U2AF35]. (C) Fold changes in SHAPE reactivity upon addition of [U2AF65 + SF1_320_ (E/E) + U2AF35] to [*AdML* + 24X U1 snRNP + 10X SRSF1-RE] (blue line) and to [*AdML* + 24X U1 snRNP + 5X SRSF1-RE] (orange line); exon-intron boundaries are demarcated with dotted vertical lines and the BS and the PPT regions are marked. (D) (Left) Amplification of *AdML* ΔPPT pre-mRNA by RT PCR from pre-cleared extract (50 µg total protein, PC), immunoprecipitant obtained from the pre-cleared extract (500 µg total protein) with anti-U2AF65 antibody (α65), and immunoprecipitant obtained from uncleared extract (500 µg total protein) with IgG2b control antibody (αCnt) performed in triplicate; (right) a plot showing significant enrichment of *AdML* ΔPPT with anti-U2AF65 antibody compared to the control antibody; error bars indicate standard deviation, ‘*’ *p*<0.05, n=3. (E) EMSA showing titration of the quaternary complexes of *β-globin* WT and *β-globin* ΔPPT assembled with SRSF1-RBD, U1 snRNP, and U2AF65 with additional U1 snRNP reduces the major soluble complex labeled as ‘ES’; the ratios of the band-intensity of the free pre-mRNA and the quaternary complexes (marked with yellow arrows inside the gel and as ‘ES’ on the side of the gel) are indicated in red script below the image; the mutated sequence in ΔPPT is shown. (F) Addition of hnRNP A1, U2AF35, and SF1_320_ (E/E) promotes PPT-dependent complexes for the WT substrate and but not the PPT mutant. (G) Summary flow chart: Cooperative binding of SRSF1 is critical for U2AF65 recruitment; however, U2AF65 non-specifically engages the pre-mRNA prior to displacement of some of the SRSF1 molecules by excess U1 snRNP; specific interactions between the pre-mRNA and U2AF65 are established upon displacement of these SRSF1 molecules; orange text indicates the steps added based on the conclusions of this Figure.

Since SRSF1 displacement is highly efficient in the presence of U2AF65, we raised the SRSF1 level from 5X to 10X molar excess to examine how the efficiency of SRSF1 displacement affects U2AF65 recruitment, (Figure 5B). After mixing 8X or 24X molar excess U1 snRNP and 50 nM [U2AF65 + SF1_320_ (E/E) + U2AF35] with the SRSF1-pre-mRNA complex, the soluble and insoluble fractions were separated (see Methods) and the MBP-MS2-bound soluble fractions were pulled down using amylose magnetic beads. We observed a similar level of U2AF65 binding to the WT and the ΔPPT variant of *AdML* in the presence of 10X SRSF1-RE. No detectable interaction with SF1_320_ (E/E) and U2AF35 was observed with either of the *AdML* variants. In contrast, in the case of 5X SRSF1-RE, specific binding of both U2AF65 as well as U2AF35 to *AdML* WT was observed (Supplementary Figure S5C). Binding of a low level of SF1_320_ (E/E) was observed to both the WT and ΔPPT variants in the absence but not the presence of U1 snRNP (lanes 2 & 5). Additionally, 24X but not 8X U1 snRNP led to a background level binding of U2AF65 to *AdML* ΔPPT substrate (Figure 5A – lanes 4-6; Supplementary Figure S5C – lane 7) likely due to partial solubilization of the precipitate (Supplementary Figure S5C – lane 13) by the equimolar arginine-HCl and glutamate-KOH (33) present in the U1 snRNP storage buffer.

To examine if the mode of binding of U2AF65 to the pre-mRNA is PPT-specific with 5X and non-specific with 10X SRSF1, we determined the SHAPE reactivity of the ternary (i.e., 1X *AdML* + *n*X SRSF1-RE + 24X U1 snRNP) and the quinary (i.e., 1X *AdML* + *n*X SRSF1-RE + 24X U1 snRNP + 50 nM U2AF65+ 50 nM U2AF35) (SF1 does not remain bound) complexes, where *n* = 5 or 10. SHAPE fold changes were calculated by dividing the SHAPE reactivity of each nucleotide of the quinary complex by that of the corresponding nucleotide of the ternary complex (Figure 5C). The plot revealed that SHAPE fold changes for complexes assembled with 5X and 10X SRSF1-RE are different across the pre-mRNA. In particular, a greater level of protection in the BS, the PPT, and the 3′SS region (165-190-nt i.e., −23 through +2 around the 3′SS) was observed in the presence of 5X SRSF1-RE compared to 10X SRSF1-RE (Figure 5C).

To examine if U2AF65 could bind the pre-mRNA in a PPT-independent manner *in vivo*, we transfected *HeLa* cells with an *AdML* ΔPPT minigene. U2AF65 was immunoprecipitated from the pre-cleared cell lysate and *AdML* ΔPPT pre-mRNA, which does not splice (8), was quantified by RT PCR (Figure 5D). Anti-U2AF65 antibody significantly enriched the pre-mRNA compared to the control antibody.

Then we examined how the titration with U1 snRNP impacts specific binding of U2AF65 to *β-globin* by EMSA since the pulldown experiment could not be done with this substrate. We added U2AF65 to the ternary complex of *β-globin* formed with 160 nM U1 snRNP and 60 nM SRSF1-RBD (Figure 5E - lanes 4-6). Addition of U2AF65 upshifted all U1 snRNP-dependent bands to a single band (labeled as ‘ES’, lane 4). Titration of the quaternary complex (ES) with increasing concentrations of U1 snRNP slightly downshifted it, likely due to displacement of additional SRSF1-RBD molecules. With *β-globin* ΔPPT, we observed that with increasing concentrations of U1 snRNP, the soluble ES complex became weaker (Figure 5E - lanes 10-12). We did not observe any detectable binding of 25 nM U2AF65 to protein-free *β-globin* pre-mRNA (Supplementary Figure S5D). Addition of SF1_320_ (E/E) and U2AF35 (Figure 5F – compare lanes 16, 18) also weakened the ES complex formed with *β-globin* ΔPPT. hnRNP A1 is reported to proofread for U2AF65 (34); accordingly, we observed a reduction in the ES complex formed with *β-globin* ΔPPT in the presence of hnRNP A1 (compare lanes 14 and 16).

Finally, we tested the assembly of the early spliceosomal ‘ES’ complex with all splice signal mutants of *β-globin*. The ES complex formed with WT *β-globin* consisted of 65% of the pre-mRNA while those formed with *β-globin* Δ5′SS, ΔBS, ΔPPT, and Δ3′SS consisted of 27%, 56%, 36%, and 64% of the mutant pre-mRNA, respectively (Supplementary Figure S5E). These results also indicate an important role of the 5’SS in the assembly of U2AF65/35. We also observed that the high efficiency of the assembly with *β-globin* ΔBS and Δ3′SS correlates with their transfection-based splicing efficiency (Supplementary Figure S5F) – *β-globin* ΔBS splices using a cryptic BS (35) and *β-globin* Δ3′SS uses a cryptic 3′SS 26-nt downstream of the authentic 3′SS.

Overall, these results suggest that excess SRSF1 bound to the pre-mRNA promotes non-specific engagement of U1 snRNP and U2AF65. Displacement of some of these SRSF1 molecules enhances the specificity of U1 snRNP and U2AF65 binding (Figure 5G). On the other hand, a lower SRSF1:pre-mRNA ratio is not sufficient to recruit U2AF65.

### Cooperation among different SR proteins for early spliceosome assembly

We sought to understand how SRSF2 and SRSF5, which also promote the splicing of *β-globin* (30), support the assembly of the early spliceosome. For this, we used the RNA binding domain (RBD) of SRSF1 (8,36,37), SRSF2-RBD in chimera with the fully phosphomimetic RS domain of SRSF1 (SRSF2-RE) (37, 38), and SRSF5-RBD (Supplementary Figure S1A). SRSF2-RE cooperatively bound *β-globin* at <3 nM concentration (Supplementary Figure S6A, lanes 1-9) while its affinity for its cognate ESE is weaker (K_d_ = ∼ 0.3 µM) (39). The complex forming the most compact band on the native gel has about 10 molecules of SRSF2-RE bound to it (6). Similarly, at least a few copies of SRSF5-RBD bound to 10 pM *β-globin* below 6 nM concentration (Supplementary Figure S6A, lane 11). Binding of SRSF5-RBD to 25 nM unlabeled *β-globin* traced with radiolabeled *β-globin* indicates that the most compact band is formed with about 15X SRSF5-RBD (Supplementary Figure S6A, lane 23). For all three SR proteins, at high SR protein to pre-mRNA probe ratios, a portion of the probe was caught in the well likely due to the formation of larger complexes that could not enter the gel.

We next tested if SRSF2 could lower the amount of SRSF1 required for cooperative engagement to *β-globin* and stable recruitment of U1 snRNP. For this, we compared the U1 snRNP-dependent *β-globin* complexes formed with varying ratios of SRSF1-RBD and SRSF2-RE. As expected, an SRSF1-RBD:*β-globin* molar ratio of less than 10 formed the *β-globin*:SR:U1 complex inefficiently (Figure 6A – lanes 3-7). However, if at least 7X molar excess SRSF2-RE was present, the lower SRSF1-RBD:*β-globin* molar ratios could assemble stable complexes (Figure 6A – lanes 8-13). To further test if the presence of multiple SR proteins helps form the *β-globin*:SR:U1 complex, we added SRSF5-RBD to the reaction mixture. In the presence of SRSF5-RBD, concentrations of both SRSF1-RBD and SRSF2-RE could be lowered to obtain similar U1 snRNP-dependent complexes (Figure 4B). Next, we examined if this cooperative relationship exists between the full-length fully phosphorylated mimetic SRSF1 and SRSF2. Although the majority of the binary and ternary complexes assembled with SRSF1-RE alone gets caught in the well under the *in vitro* experimental conditions (Supplementary Figure S6B, lanes 7-11), lower concentrations of SRSF1-RE in the presence of a low concentration of SRSF2-RE formed U1 snRNP-dependent complexes similar to SRSF1-RBD (Supplementary Figure S6B, lanes 13-18).

**Figure 6.**
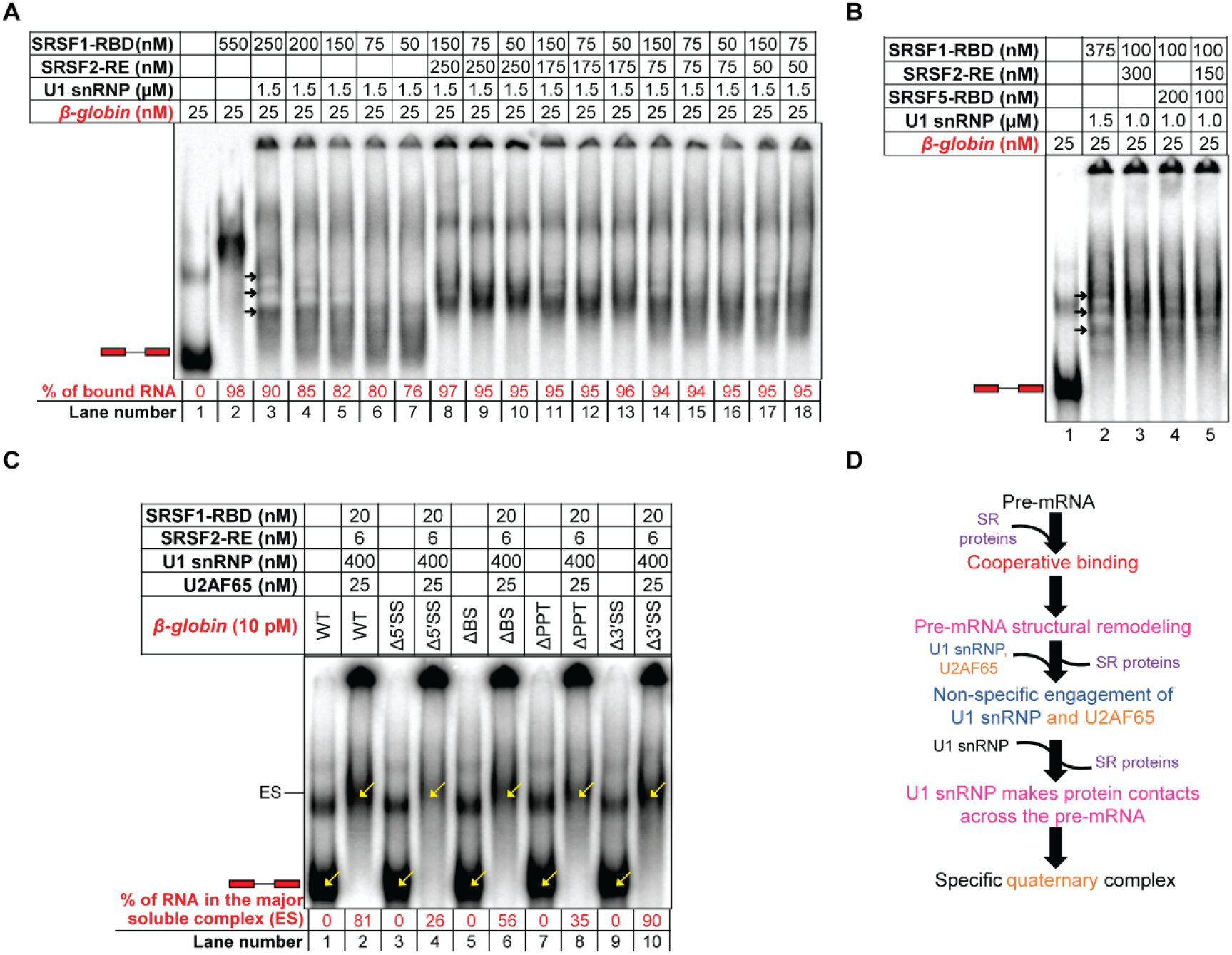
Collaboration of SR proteins for assembly of the early spliceosome. (A) EMSA showing U1 snRNP stabilization efficiency of 10X, 8X, 6X, 3X, and 2X SRSF1-RBD on 25 nM *β-globin* (lanes 4-7), with 6X, 3X, and 2X SRSF1-RBD and 10X SRSF2-RE (lanes 8-10), with 6X, 3X, and 2X SRSF1-RBD and 7X SRSF2-RE (lanes 11-13), with 6X, 3X, and 2X SRSF1-RBD and 3X SRSF2-RE (lanes 14-16), with 6X and 3X SRSF1-RBD and 2X SRSF2-RE (lanes 17-18). (B) EMSA showing formation of comparable stable *β-globin*:SR:U1 snRNP complexes in the presence of a high level of SRSF1-RBD alone or a low level of SRSF1-RBD supplemented with SRSF2-RE and/or SRSF5-RBD. (C) EMSA showing a high efficiency of assembly of the early spliceosomal complexes with U1 snRNP and U2AF65 in the presence of low concentrations of SRSF1-RBD and SRSF2-RE with the splicing-competent variants of *β-globin* (WT, ΔBS, Δ3′SS). (D) Summary flow chart: A combination of different pre-mRNA-specific SR proteins at low SR:pre-mRNA ratios could substitute for a high SRSF1:pre-mRNA ratio required for assembly of the early spliceosomal complexes; violet text indicates the steps added based on the conclusions of this Figure.

Then we examined U2AF65-dependent complex formation with a combination of [6 nM SRSF2-RE + 20 nM SRSF1-RBD] instead of 60 nM SRSF1-RBD alone (Figure 6C) using *β-globin* and its splice signal mutants. The major soluble complex was most intense for the splicing competent variants (WT, ΔBS, and Δ3′SS – lanes 2, 6, and 10). These results suggest that the SR proteins collaborate to recruit early spliceosome factors lowering each other’s concentration requirement and that all splice signals are important for the recruitment of U2AF65.

Overall, these results suggest that the molar ratio of individual SR proteins to the pre-mRNA dictates early spliceosome assembly efficiency (Figure 6D).

## DISCUSSION

The mechanisms governing the recognition of the highly degenerate mammalian splice signals by the early spliceosomal components from background sequences remain unclear. Data shown in this and our earlier work (8) suggest that pre-mRNAs adopt a folded structure integrating all splice signals and such early spliceosomal components as U1 snRNP, U2AF65, and U2AF35 recognize the 3D structural scaffold. However, the scaffold needs to be remodeled for recognition and/or stabilization of the early spliceosomal components. Data shown in the current work suggest that cooperative binding of SR proteins to the pre-mRNA enables this structural remodeling. The cooperative nature of binding also explains how SR proteins bind the pre-mRNA efficiently *in vivo* despite having a weak affinity (in the micromolar range) for their weakly conserved binding motifs. This cooperative binding requires a minimum number of SR protein molecules per molecule of the pre-mRNA. Interestingly, pre-mRNA-bound SRSF1 at a higher SRSF1:pre-mRNA ratio promotes non-specific engagement of U1 snRNP and U2AF65 to the tested splicing substrates. We find that a large excess of U1 snRNP, which is overabundant in living cells compared to other U snRNPs (40), selectively displaces some SRSF1 molecules from the pre-mRNAs enabling the splice signal-specific engagement of U1 snRNP, U2AF65, and U2AF35. These data also explain why assembled early spliceosome contains only a few SR protein molecules bound to it (20) despite having SR protein binding motifs scattered across the entire pre-mRNA. This model of splicing substrate recognition is depicted in a cartoon in Figure 7. The SRSF1-dependent non-specific U1 snRNP engagement was easily detected with the *AdML* ΔPPT mutant, which is completely defective in U1 snRNP recruitment in the protein-free state due to a disrupted scaffold (8). However, it was not possible to examine this phenomenon with *β-globin* because its 5′SS-mutant in the protein-free state is highly similar to the WT substrate in its structure and the ability to recruit U1 snRNP. Nonetheless, these observations tempt us to propose that the non-specific engagement of splicing factors prior to displacement of SRSF1 (or other SR proteins) increases the probability of specific engagement of the former after displacement of the latter, thus likely improving early spliceosome assembly efficiency. Additionally, the SR protein displacement step has the potential to act as a regulatory checkpoint for early spliceosome assembly *in vivo*. This hypothesis is consistent with the earlier observations that an increase in SR protein levels could disrupt efficient early spliceosome assembly and splicing (19,41,42).

**Figure 7.**
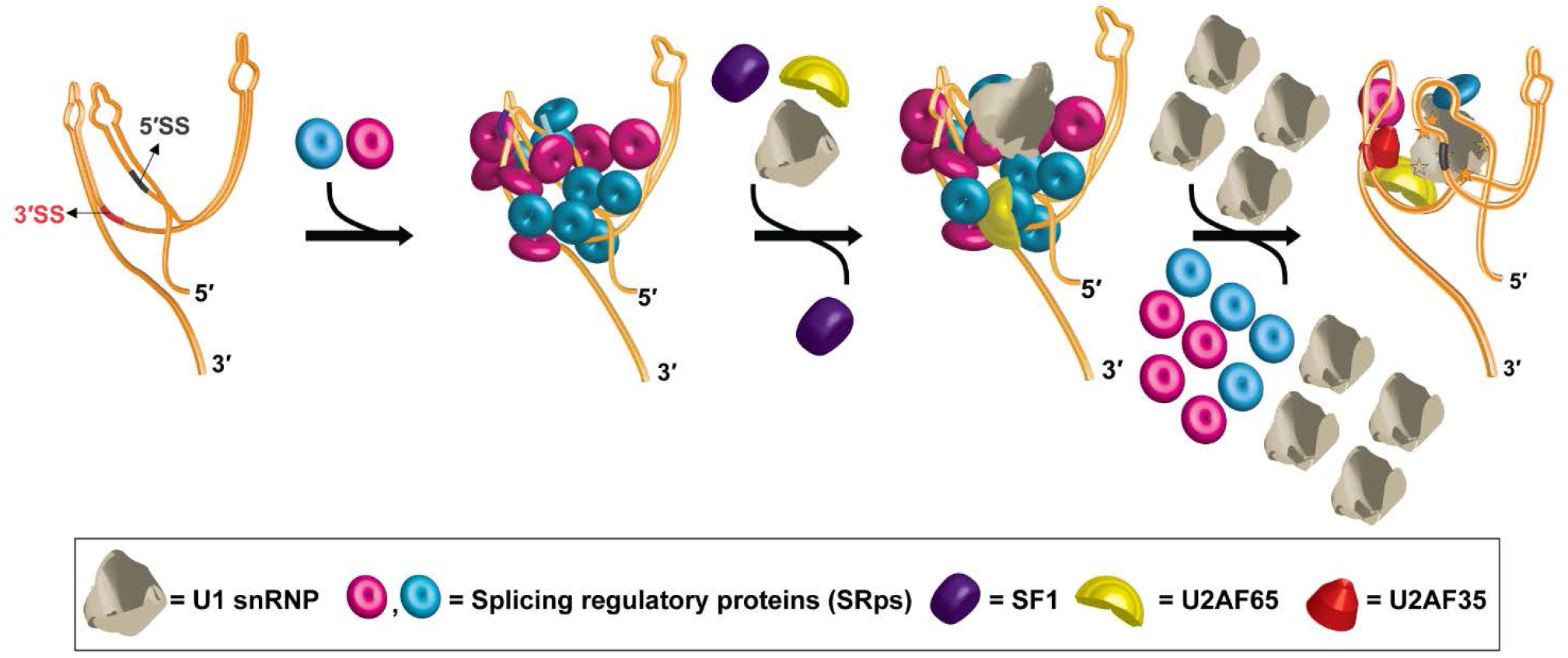
Proposed model for mammalian splicing substrate definition by cooperative binding and subsequent selective displacement of SR proteins. The primary, secondary, and tertiary structure of a pre-mRNA mediate the cooperative recruitment of multiple copies of different SR proteins (and potentially other splicing regulatory proteins) specific to the pre-mRNA. The SR:pre-mRNA complex initially recruits U1 snRNP and U2AF65 non-specifically. A large excess of U1 snRNP selectively displaces some SR protein molecules from the pre-mRNA leading to structural remodeling of the pre-mRNA and specific recruitment of U1 snRNP, U2AF65, and U2AF35. SF1, which appeared to be critical for recruitment of U2AF65 and U2AF35, is detected in the pre-mRNA complex prior to displacement of the majority of the SR protein molecules. However, it is not detected in the same pre-mRNA complex consisting of U2AF65, U2AF35, and U1 snRNP suggesting its chaperone-like role as proposed earlier (8).

U1 snRNP-mediated SR protein displacement appears to involve a complex and dynamic mechanism. SRSF1 interacts with U1-70k (37) and the stem-loop III of U1 snRNA (43). On the other hand, U1 snRNP interacts with various segments of the pre-mRNA [(8) and Figure 4D]. Therefore, we propose that a combination of interactions between U1 snRNP and SRSF1 and between U1 snRNP and the pre-mRNA are primarily responsible for U1 snRNP-mediated SRSF1 displacement. Interestingly, U1 snRNP B_174_ ΔA displaces SRSF1 more efficiently than U1 snRNP B_174_ despite having a weaker interaction with the pre-mRNA and SRSF1 compared to U1 snRNP; we hypothesize that transient interactions of this variant with the pre-mRNA and SRSF1 help recycle this variant more efficiently. Further studies are required to examine this possibility. Displacement of other SR proteins by U1 snRNP may also involve similar mechanisms, many of which exhibit interactions with U1 snRNP (43). The selective displacement of SRSF1 (or other SR proteins) from the pre-mRNA is likely guided by the position of its binding site within the 3D scaffold and the affinity of the site for the protein. After displacement, the remaining SRSF1 bound to the pre-mRNA complex could be present within the early spliceosome by bridging the SRE and U1 snRNP (20) or by bridging U1 snRNP and another early spliceosomal component at the 3′ end of the intron (43). Currently it is not clear which mode is being used for the tested splicing substrates. Nonetheless, SRSF1 bound to the pre-mRNA:U1 snRNP:SRSF1 complex are ‘locked’ within the pre-mRNA structural scaffold and are not displaced at least until after the recruitment of U2AF65 and U2AF35.

In this study, a truncated U1-70k (1-215 a.a.) is used to assemble U1 snRNP. At least two different isoforms of U1-70k participate in the assembly of human U1 snRNP. These two isoforms, which differ in amino acid sequence beyond the 215^th^ residue, exhibit different interactions within the U1 snRNP particle and different phosphorylation patterns (44). Without additional knowledge about their functionalities, it is not possible to ascertain whether and how selective usage of the one or the other full-length isoform or the truncated protein impacts the results. However, the residues beyond the 215^th^ position of U1-70k constitutes the RS and R-D/E domain, which primarily serves as a regulator of the functions of U1-70k for assembly of U1 snRNP; this domain is phosphorylated *in vivo* for the release of the RNA recognition motif (RRM) from the contact with the RS+R-D/E domain, an essential step for assembly of U1 snRNP (45). That is, in the absence of the RS+R-D/E domain, U1-70k may be constitutively active. An earlier finding that the RS domain of U1-70k is dispensable for the survival of simpler organisms such as *Drosophila* supports this hypothesis (46). Given the situation, we surmise that the observations made with U1 snRNP assembled with the truncated U1-70k would resemble the behavior of native U1 snRNP *in vivo*. Nonetheless, the physiological and/or clinical relevance of these observations remains to be established.

Data presented in this and our earlier manuscript (8) bring to light a novel regulatory niche in the early spliceosome assembly by providing insights into the regulation of early spliceosome assembly in the context of the pre-mRNA 3D structural scaffold. We observed several common as well as different features in the tested viral and human transcripts allowing for diversified splicing regulation. For example, although *AdML* 5′SS has six nucleotides complementary to the 5′ end of U1 snRNA while *β-globin* 5′SS has seven, the *AdML* scaffold binds U1 snRNP more stably with a greater involvement of its 5′SS than the *β-globin* scaffold. On the other hand, both *AdML* and *β-globin* scaffolds are similar in requiring the 5′SS for U2AF65 recruitment, suggesting cross-intron communication through the 3D space in both substrates. Besides, although SF1 is known to regulate splicing genome-wide (47), its mechanisms of action remain unclear (48). Our data obtained with a truncated variant of SF1 containing the N-terminal 320 amino acids suggest that SF1 but not U2AF65 and U2AF35 remains bound to the pre-mRNA:SRSF1 complex at low U1 snRNP concentrations, but with increasing U1 snRNP concentrations, U2AF65 and U2AF35 binding improves and the SF1 level diminishes. This is consistent with our earlier hypothesis that SF1 might have a chaperone-like function for the recruitment of U2AF65 and U2AF35 (8). Finally, the inability of individual binding motifs to explain the splicing outcome and the cooperative binding are also reported for many other SRps (49–55). Thus, we infer that the mechanistic knowledge presented in the current work using SRSF1 could also be extended to a variety of other SRps.

Overall, this work highlights the highly complex mode of early spliceosome assembly where SR proteins endow a greater flexibility to the pre-mRNA 3D structural scaffold. U1 snRNP facilitates its own recruitment and recruitment of U2AF65 and U2AF35 by selectively removing some SR protein molecules. Despite generation of several high-resolution structures for various stages of the spliceosome in the recent years, both the structure and the mechanism of assembly of the early spliceosome, particularly E-complex, are still unclear (56). Our data expand on the novel regulatory niche in the early spliceosome assembly by characterizing the detailed mechanism of modulation of the pre-mRNA 3D structural scaffold and assembly of the mammalian E-complex, thus paving the way for its detailed structural analysis. Additionally, the knowledge of pre-mRNA-wide dynamic binding of RBPs generated in this work will be valuable in developing RNA therapeutics (57) targeting previously unthought of segments of a pre-mRNA to alter the RBP binding and/or the 3D scaffold in order to rescue splicing defects.

## AUTHOR CONTRIBUTION

KS designed and performed the experiments, analyzed the data, and wrote the paper. GG designed some experiments, wrote the paper, and supervised the project.

## ACKNOWLEDGEMENT

The authors would like to acknowledge Dr Simpson Joseph for critical reading of the manuscript and Mike Minh Fernandez for some early work.

## DATA AVAILABILITY

Raw and processed files pertaining to the RNA-seq data have been deposited to Gene Expression Omnibus under accession number GSE188652.

