## Supplementary Figures for "Cooperative engagement and subsequent selective displacement of SR proteins define the pre-mRNA 3D structural scaffold for early spliceosome assembly"

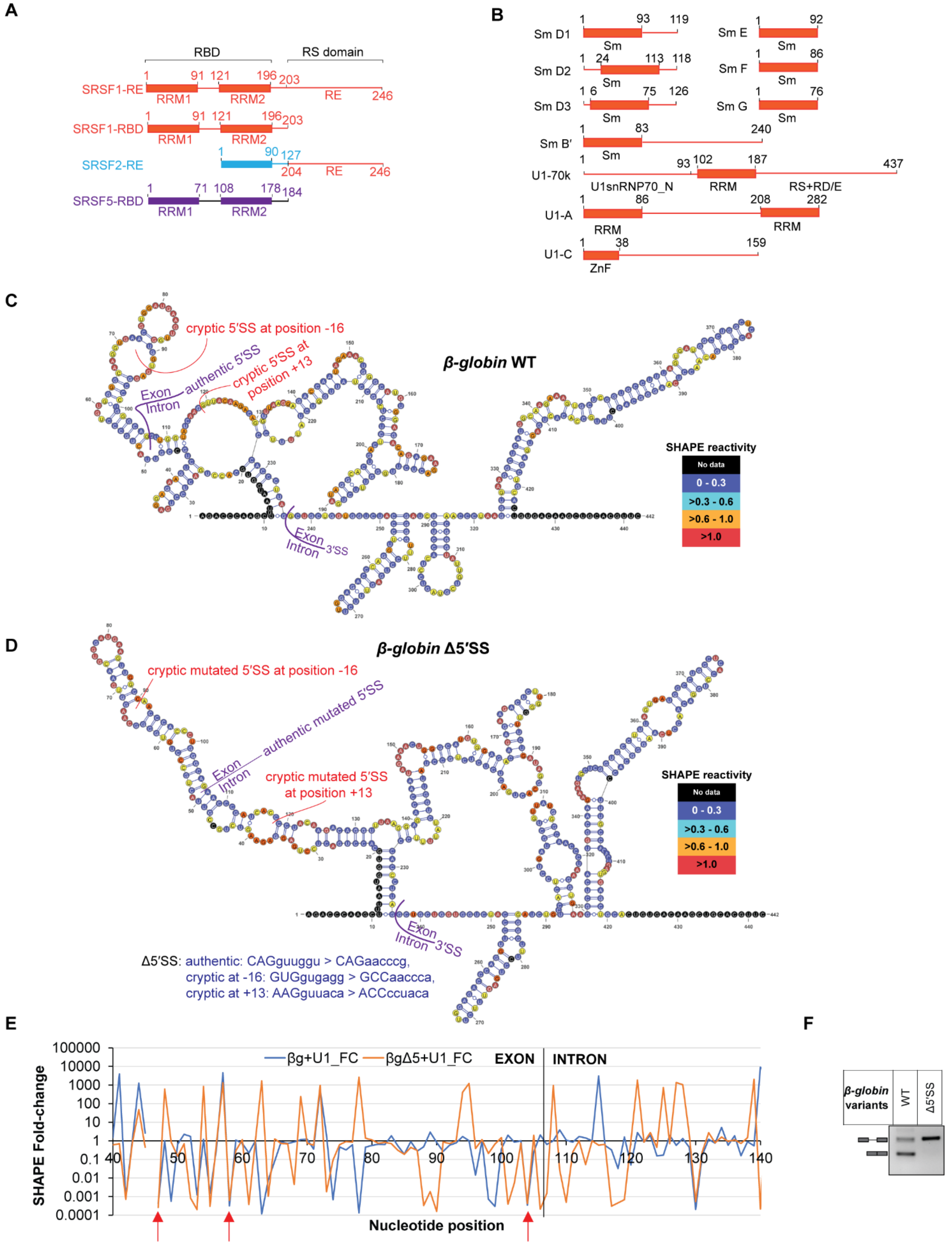
A
B
C
D
E
F
RBD
RS domain
SRSF1-RE
SRSF1-RBD
SRSF2-RE
SRSF5-RBD
RRM1
RRM2
RE
Sm D1
Sm D2
Sm D3
Sm B′
U1-70k
U1-A
U1-C
Sm E
Sm F
Sm G
Sm
U1snRNP70_N
RRM
RS+RD/E
ZnF
β-globin WT
cryptic 5′SS at position -16
authentic 5′SS
cryptic 5′SS at position +13
Exon
Intron
3′SS
SHAPE reactivity
No data
0 - 0.3
>0.3 - 0.6
>0.6 - 1.0
>1.0
β-globin Δ5′SS
cryptic mutated 5′SS at position -16
authentic mutated 5′SS
cryptic mutated 5′SS at position +13
Δ5′SS: authentic: CAGguuggu > CAGaacccg,
cryptic at -16: GUGgugagg > GCCaaccca,
cryptic at +13: AAGguuaca > ACCccuaca
βg+U1_FC
βgΔ5+U1_FC
EXON
INTRON
SHAPE Fold-change
Nucleotide position
β-globin variants
WT
Δ5′SS

**Supplementary Figure S1. Global structure of *β-globin* variants and its interactions with U1 snRNP**

(A) SR protein constructs used in this study. (B) Domain architecture of U1 snRNP component proteins. (C & D) SHAPE-derived secondary structure models of *β-globin* WT (C) and its 5′SS mutant (D); the sequences mutated in *β-globin* Δ5′SS are shown; nucleotides are color-coded according to their SHAPE reactivity as indicated in the associated table. (E) Fold changes in SHAPE reactivity in the segments flanking the 5′SS upon U1 snRNP engagement to protein-free *β-globin* and *β-globin* Δ5′SS. (F) Transfection-based splicing assay showing the lack of splicing ability of *β-globin* Δ5′SS mutant.

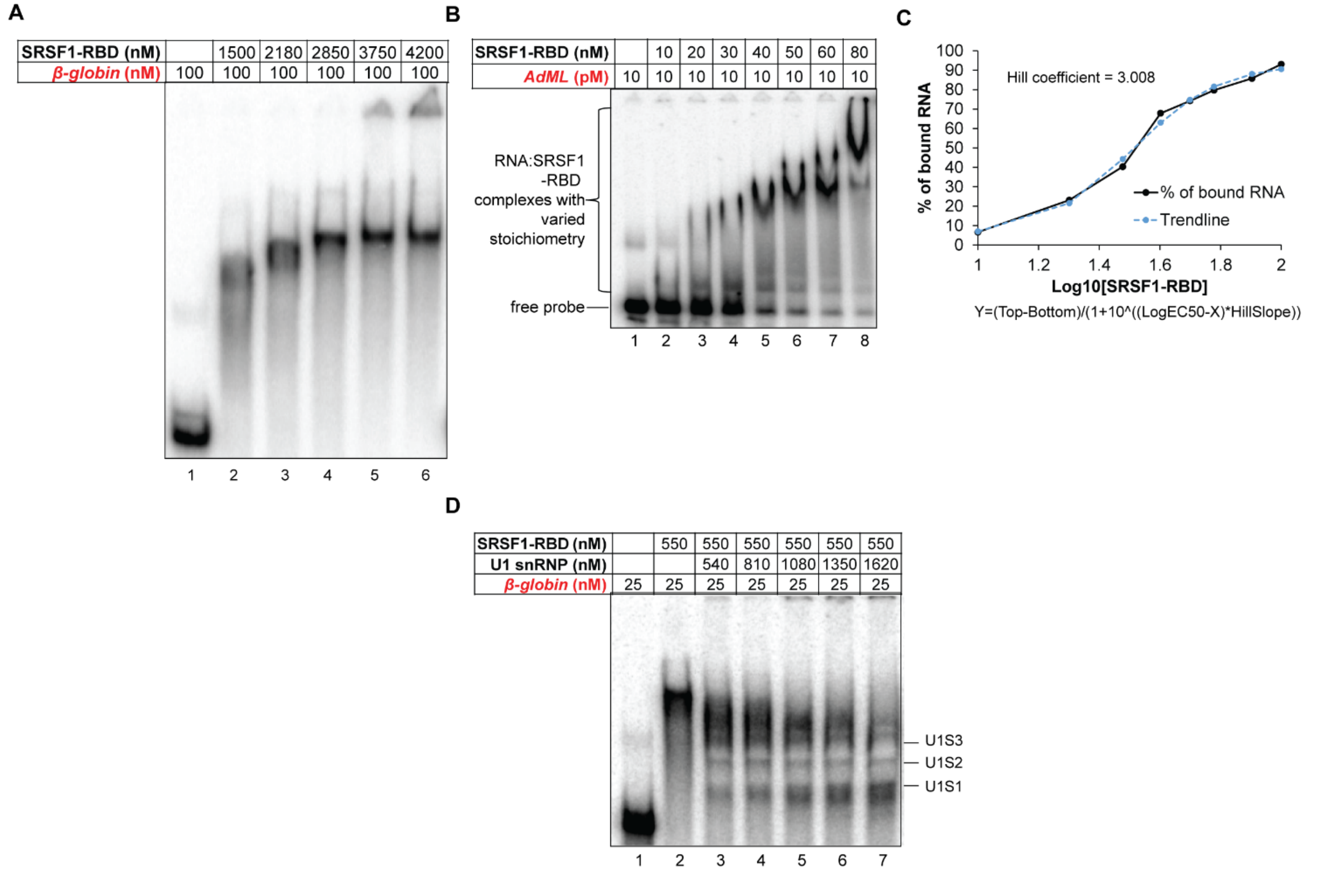


**Supplementary Figure S2. Cooperative SRSF1-RBD binding and U1 snRNP recruitment to pre-mRNAs**

(A) Titration of 100 nM *β-globin* traced with radiolabeled *β-globin* with SRSF1-RBD showing binding of multiple copies of SRSF1-RBD to each *β-globin* molecule; radiolabeled components are shown in red script. (B) Cooperative binding of SRSF1-RBD to 10 pM radiolabeled *AdML* pre-mRNA. (C) Calculation of Hill coefficient from the band intensity of the free probe in B; the equation of the binding curve (solid black line) is given below it, where Y is band intensity, X is $Log_{10}$[SRSF1], and top and Bottom are plateaus in the unit of the Y axis; a trendline (broken blue line) was generated to visually examine the goodness of fit by estimating the Y values from the X-values, Hill Slope (3.008), EC50 (32.17), Top (93.4), and Bottom (4.373). (D) About 60X molar excess U1 snRNP is required to resolve the U1 snRNP-dependent ternary complexes formed with 25 nM *β-globin* bound to 22X molar excess SRSF1-RBD.

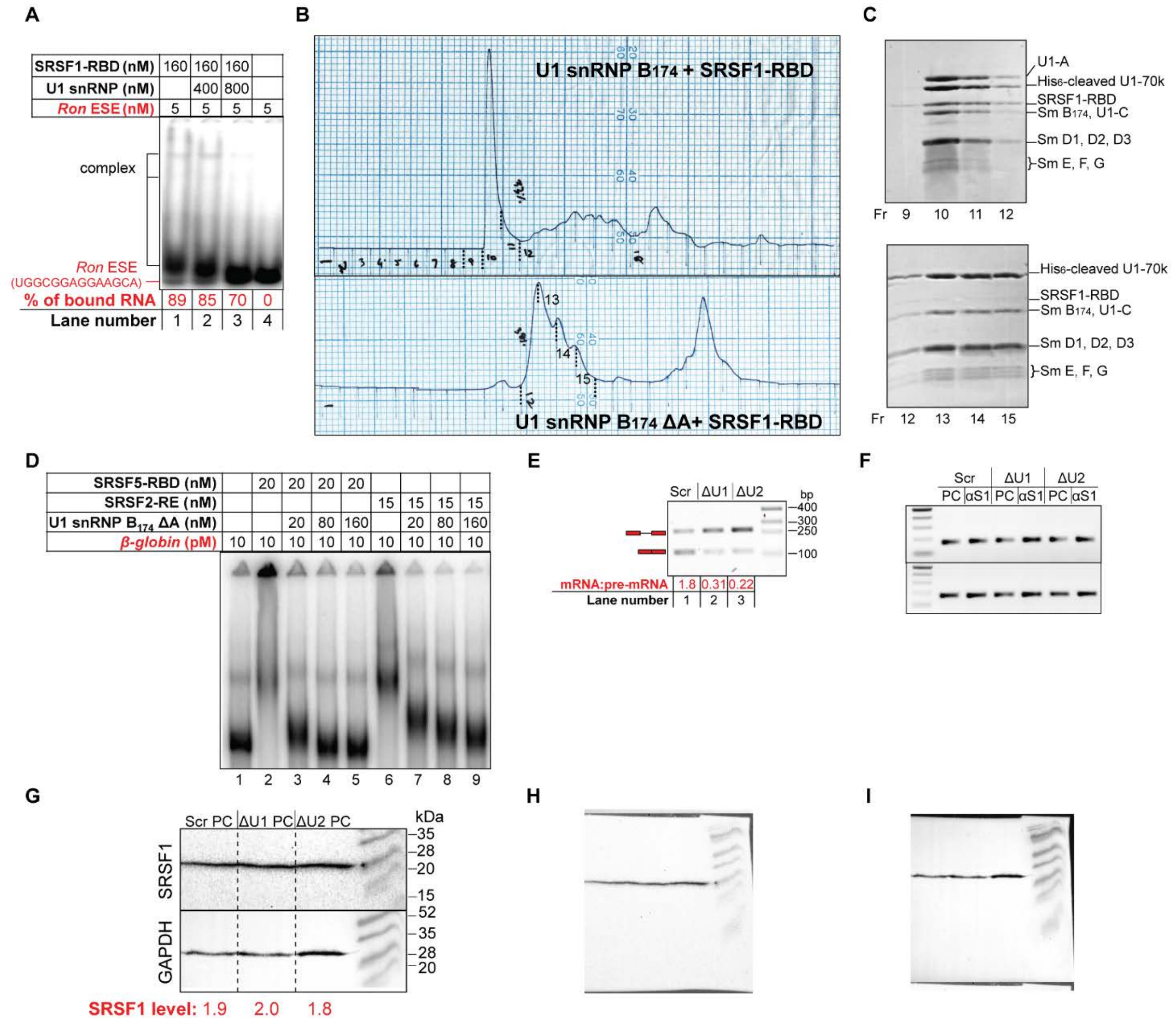


**Supplementary Figure S3. Displacement of SRSF1-RBD from the RNA by U1 snRNP**

(A) Titration of *Ron* ESE-SRSF1-RBD complexes with U1 snRNP showing displacement of SRSF1-RBD from the RNA releasing free probe. (B) Chromatograms of purification of complexes formed with U1 snRNP $B_{174}$ and SRSF1-RBD (top) and U1 snRNP $B_{174}$ ΔA and SRSF1-RBD (bottom) by anion-exchange chromatography. (C) SDS PAGE of peak fractions showing formation of 1:1 U1 snRNP $B_{174}$-SRSF1-RBD complex (top) but no detectable complex between U1 snRNP $B_{174}$ ΔA and SRSF1-RBD (bottom). (D) EMSA showing displacement of SRSF2-RE and SRSF5-RBD from *β-globin* by U1 snRNP $B_{174}$ ΔA. (E) Transfection-based splicing assay showing reduction in splicing efficiency of *AdML* pre-mRNA upon treatment of *HeLa* cells with U1 AMO (ΔU1) and U2 AMO (ΔU2) compared to scrambled AMO (Scr). (F) Two replicates of the result shown in Figure 3F. (G) Detection of SRSF1 levels by western blot in cells treated with scrambled, U1, and U2 AMO; normalized SRSF1 band intensity based on GAPDH internal standard is shown below. (H & I) Images of the full blot probed with anti-SRSF1 antibody (H) and anti-GAPDH antibody (I).

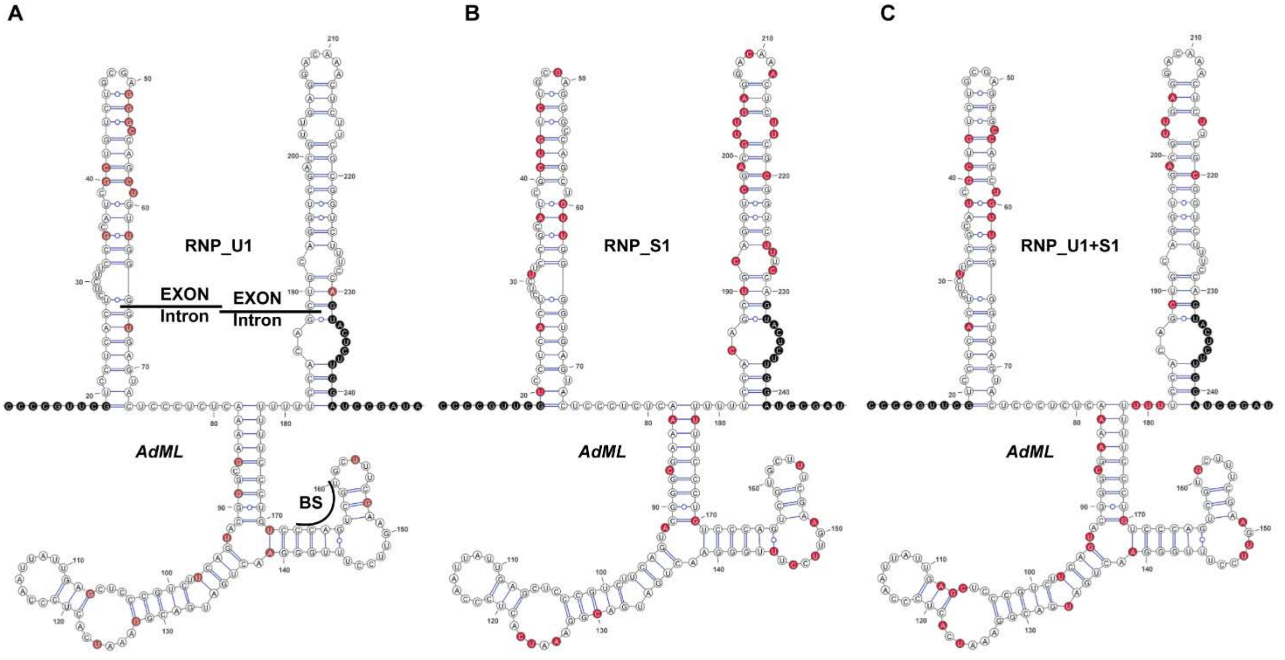


**Supplementary Figure S4. *In vitro* RNP-reactivity of *AdML***

(A, B, C) RNP-MaP-derived lysine-contacting nucleotides in U1 snRNP- (A), SRSF1-RE (B), or U1 snRNP+SRSF1-RE- (C) bound *AdML* are marked in red in the SHAPE-derived secondary structure model of *AdML*.

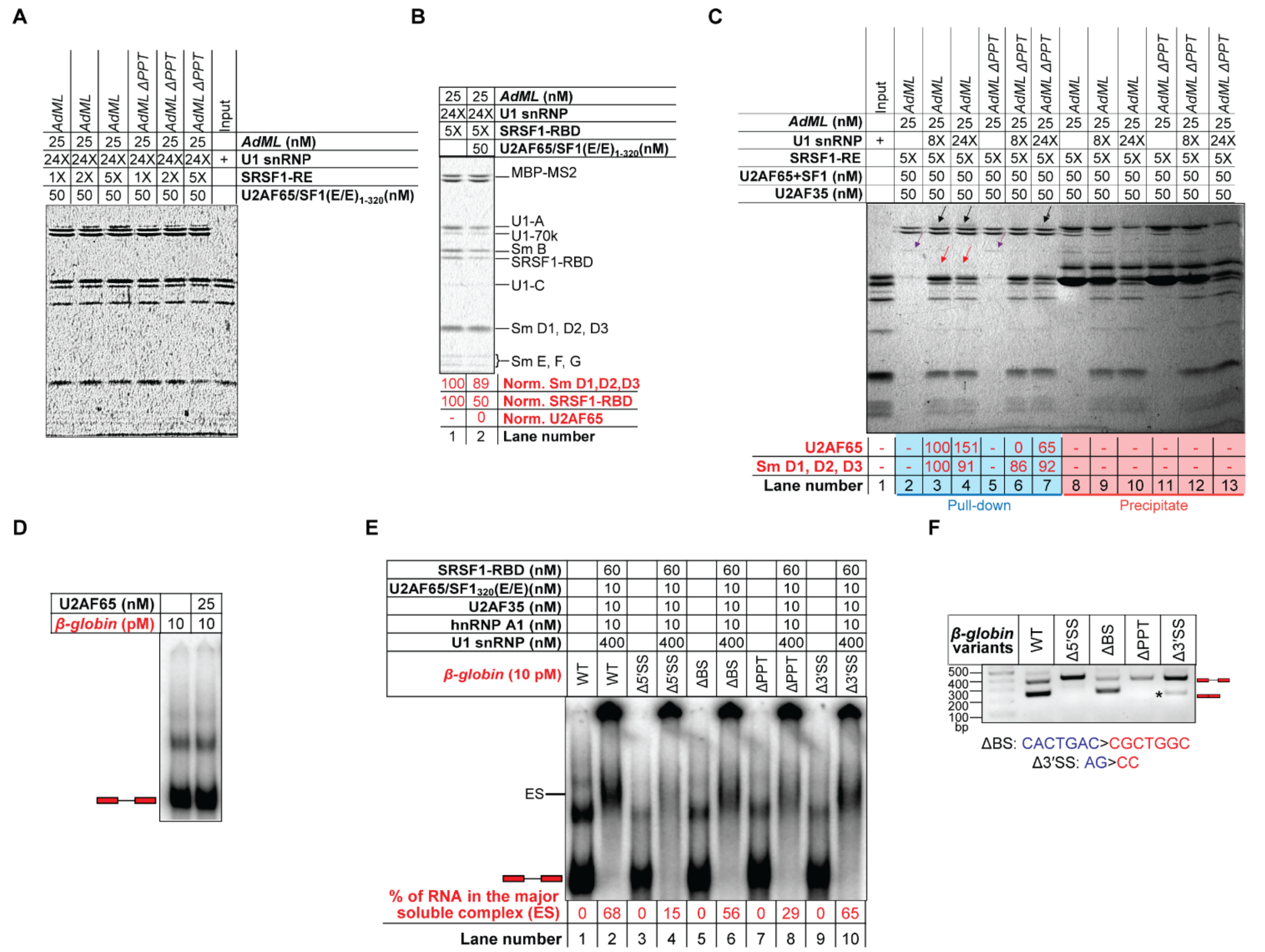

**Supplementary Figure S5. U2AF65 recruitment requires engagement of SRSF1 in appropriate stoichiometry and then its displacement by U1 snRNP**

(A) Background-subtracted and contrasted version of Figure 5A. (B) Amylose pull-down assay of MS2-tagged *AdML* in the presence of 5X SRSF1-RBD, 24X U1 snRNP, and 0 or 50 nM [U2AF65 + $SF1_{320}$ (E/E)]. (C) Amylose pull-down assay of MS2-tagged *AdML* in the presence of 8X or 24X molar excess U1 snRNP, 50 nM [U2AF65 + $SF1_{320}$ (E/E) + U2AF35], and 5X SRSF1-RE; black arrows indicate U2AF65, red arrows U2AF35, and violet arrows $SF1_{320}$ (E/E). (D) U2AF65, at 25 nM concentration, does not exhibit a detectable binding to full-length protein-free *β-globin*. (E) EMSA showing a high efficiency of assembly of the early spliceosomal complex labeled as 'ES' with SRSF1-RBD, U2AF65, $SF1_{320}$ (E/E), and hnRNP A1 for the splicing-competent variants of *β-globin* (WT, ΔBS, Δ3′SS). (F) Transfection-based splicing assay of *β-globin* WT and splice signal mutants; '*' indicates an mRNA produced using a cryptic 3′SS 26-nt downstream of the authentic 3′SS.

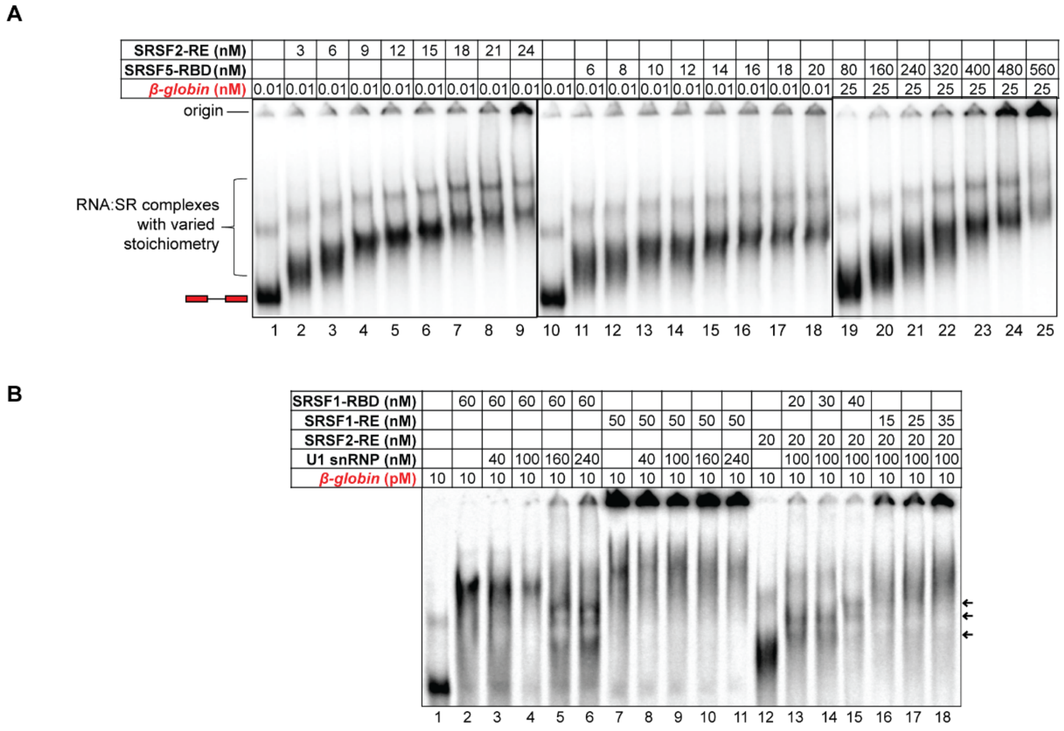


**Supplementary Figure S6. Cooperative actions of different SR proteins**

(A) EMSA showing titration of 10 pM *β-globin* with SRSF2-RE (lanes 1-9) and SRSF5-RBD (lanes 10-18), and 25 nM *β-globin* with high concentrations of SRSF5-RBD (lanes 19-25). (B) EMSA showing U1 snRNP-dependent complex formation with *β-globin* in the presence of SRSF1-RBD alone (lanes 2-6), SRSF1-RE alone (lanes 7-11), low concentrations of SRSF1-RBD and SRSF2-RE (lanes 13-15), and low concentrations of SRSF1-RE and SRSF2-RE (lanes 16-18).
